# Temporal blood sampling improves parameter recovery in a model of enterohepatic bile acid circulation using surrogate-accelerated Bayesian inference

**DOI:** 10.64898/2026.11.18.766999

**Authors:** Sebastian Joseph, Isabelle A. Leclercq, Laure-Alix Clerbaux, Sophie de Buyl

## Abstract

Bile acids (BAs) are essential signaling molecules whose enterohepatic circulation (EHC) is disrupted across hepatobiliary disorders. Despite their potential as biomarkers, clinical utility is limited by high inter-individual variability, the inability of accessible peripheral blood to reflect true hepatic exposure, and the difficulty of directly measuring key physiological parameters governing the EHC. While mechanistic models can infer inaccessible hepatic profiles and patient-specific parameters from peripheral blood, personalizing them is computationally prohibitive due to multiscale dynamics causing numerical stiffness and the thousands of evaluations required for Bayesian inference.

To overcome this, we developed an accelerated Bayesian inference pipeline combining timescale model reduction with neural network surrogates. Eigenvector participation factor analysis of a human EHC model revealed that rapid portal vein and hepatic sinusoidal transport drive stiffness. Applying the quasi-steady state approximation to these fast variables reduced integration times by over 97% with an integrated relative error <1.0%. A neural network surrogate further accelerated population ensemble Markov chain Monte Carlo inference from days to hours, accurately recovering parameter distributions in 100 virtual individuals (95-98% coverage).

Crucially, this tractability allowed us to dissect EHC physiological mechanisms and their disease perturbation. Single fasting measurements yielded practical non-identifiability due to strong posterior correlations among hepatic synthesis, intestinal motility, and absorption. A systematic comparison of sampling protocols showed peripheral sampling at multiple time points progressively disentangles these processes, while portal vein sampling offers no added benefit. Three peripheral time points emerged as an optimal protocol, recovering >90% of the identifiability of five time points. Furthermore, the framework captured EHC compensatory dynamics, revealing a persistent positive correlation between intestinal transit and absorption.

By making stiff EHC models tractable, our framework guides clinical study design and enables minimally invasive, mechanistic characterization of BA dysregulation, advancing our mechanistic understanding and personalized diagnosis of hepatobiliary diseases.

## Introduction

### Bile acids as promising biomarkers for liver diseases

Besides their well described role in dietary fats digestion, bile acids (BAs) are recognized as key metabolic signaling molecules. By binding to cognate receptors, they regulate hepatic inflammation as well as glucose and lipid homeostasis [1, 2]. BAs are synthesized in the liver to form primary BAs that are then conjugated to taurine or glycine. After secretion into the intestine via the bile, they are transformed to secondary BAs by the gut microbiota. The vast majority (>95%) are reabsorbed into the portal circulation, and taken back into the liver. Only a small proportion of BAs are excreted through the feces or leak into the peripheral circulation. Hence, BAs are continuously recycled through this so-called enterohepatic circulation (EHC) (Fig 1), a highly dynamic and tightly regulated process. BA synthesis is controlled by feedback from both hepatic BAs and intestinal BAs through the Farnesoid X Receptor-Fibroblast Growth Factor 19 (FXR–FGF19) axis, while meals trigger gallbladder emptying and modulate intestinal transit. BA homeostasis further depends on intestinal absorption, microbial transformation, and hepatic uptake. Together, these processes determine the size and composition of the circulating BA pool.

**Figure 1.**
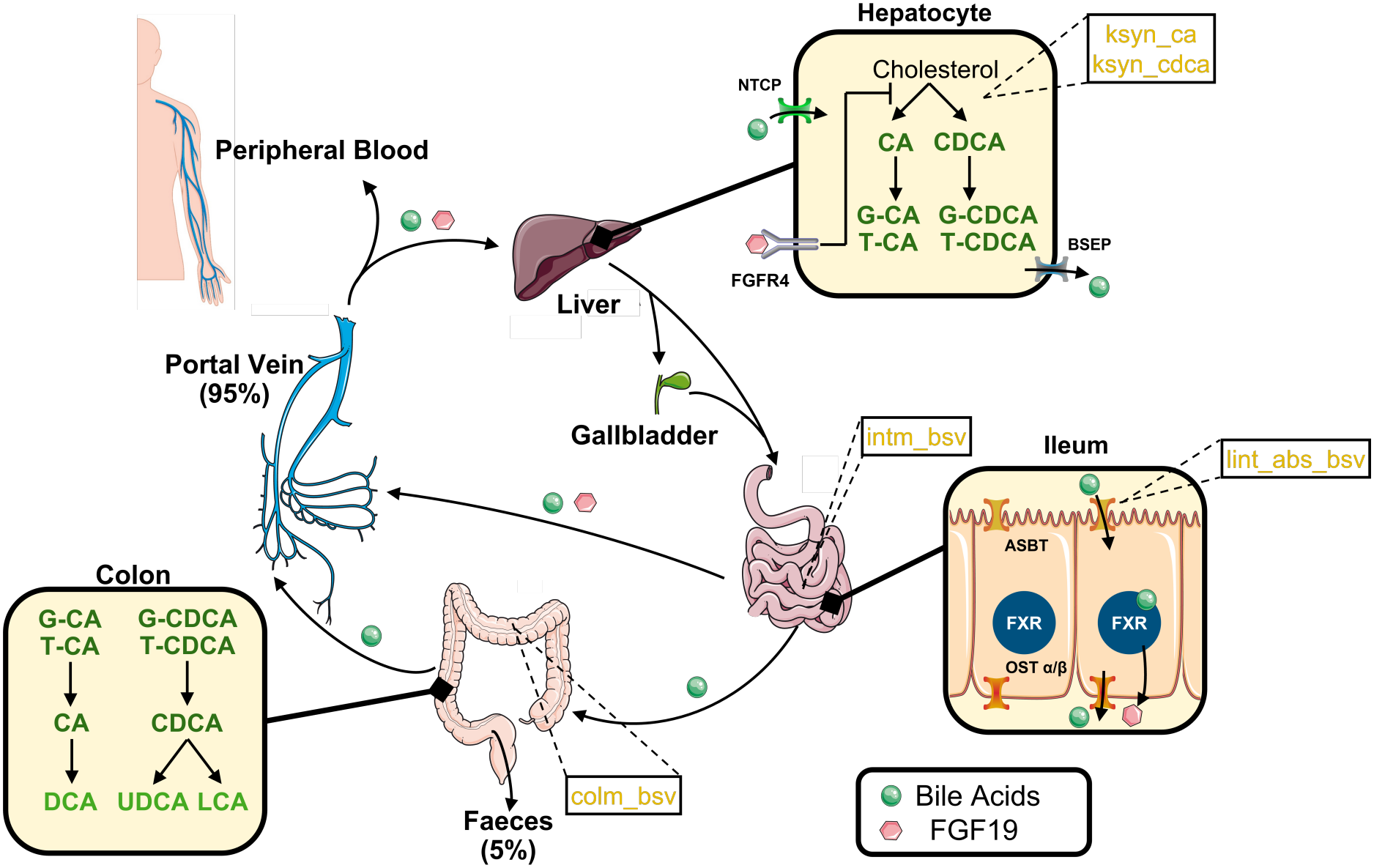
Schematic of the human enterohepatic circulation. Adapted from the enterohepatic circulation of bile acids described in Joseph *et al*. [3]. The five inferred model parameters in this study are in orange and boxed (e.g. `ksyn_ca`). *Abbreviations: ASBT, Apical Sodium-Bile acid Transporter; BSEP, Bile Salt Export Pump; CA, cholic acid; CDCA, chenodeoxycholic acid;* `colm_bsv`, *colonic motility rate scalar; DCA, deoxycholic acid; FGF19, Fibroblast Growth Factor 19; FGFR4, Fibroblast Growth Factor Receptor 4; FXR, Farnesoid X Receptor; G-, glycine-conjugated;* intm*_bsv*, *intestinal motility rate scalar;* `ksyn_ca`, *synthesis rate of CA;* `ksyn_cdca`, *synthesis rate of CDCA; LCA, lithocholic acid;* `lint_abs_bsv`, *lower intestinal absorption rate scalar; NTCP, Na*^+^*-Taurocholate Cotransporting Polypeptide; OSTα*/*β, Organic Solute Transporter alpha/beta; T-, taurine-conjugated; UDCA, ursodeoxycholic acid*.

Disruption of the EHC of BAs has been implicated in a range of hepatobiliary disorders, including metabolic dysfunction-associated steatotic liver disease (MASLD), cholestasis and drug-induced liver injury [3, 4, 5]. These conditions represent a substantially growing clinical burden, with MASLD affecting approximately one-third of the global adult population [6]. MASLD can progress to more fatal conditions such as cirrhosis and hepatocellular carcinoma, while cholestasis and drug-induced liver injury are leading causes of acute liver failure [7, 8]. However, despite more than three decades of research, the use of peripheral BAs as biomarkers for human liver injury remains inconclusive due to inconsistent findings [3]. While some studies report strong correlations between elevated BA levels and disease severity, this mainly pertains to end-stage liver disease, where liver function is compromised and portal hypertension with vascular shunting has developed. In contrast, other studies find weak or no associations, and results vary depending on whether total BA concentrations or individual BA species are measured, as well as whether samples are collected under fasting or postprandial conditions. Besides the need to harmonize methodological techniques between studies, two major challenges have limited the clinical relevance of BAs as biomarkers so far.

Firstly, intra-individual portal and peripheral BA profiles differ markedly [9], meaning that clinically accessible peripheral blood cannot reliably predict hepatic exposure to portal BAs. Yet, comprehensive characterization of portal BAs in humans is severely limited by logistical and ethical constraints. A mathematical model of the entire EHC is an invaluable tool to infer the portal BA profiles from accessible clinical measurements.

Secondly, peripheral BA concentrations show very high inter-individual variability, arising from variations throughout the EHC, for example variations in hepatic BA synthesis or in gut microbiota composition affecting colonic transformation towards secondary BAs. Tailoring the EHC of BAs to each patient or to a subgroup of patients is a prerequisite towards personalized diagnoses. However, the multi-organ complexity of the EHC makes it impossible to predict experimentally how inter-individual variations of each parameter affects the BA pool throughout the EHC. Here again, a mathematical model can leverage clinical measurements with physiological knowledge, linking changes in specific processes, such as biliary secretion or microbial metabolism, to BA profiles.

### Mechanistic models for physiological insight and parameter inference

Mechanistic models of biological systems are constructed by taking into account our current knowledge of the underlying physiological mechanisms. They are particularly valuable since they capture the intrinsic dynamic nature of living systems, i.e. how they constantly evolve over time. Analysis of these models often provide insights into interactions between variables and parameters that are difficult to investigate experimentally, such as determining individual BA hepatic uptake rates or ileal reabsorption. Mechanistic models can infer these inaccessible physiological parameters *in silico* by fitting the model to minimally invasive clinical measurements, such as peripheral blood and fecal BA concentrations.

A detailed model of the human EHC has been constructed by Voronova *et al*. [10]. The model consists of 74 ordinary differential equations (ODEs) capturing the dynamics of cholic acid (CA), chenodeoxy-cholic acid (CDCA), deoxycholic acid (DCA), and their taurine and glycine conjugates (T-CA, G-CA, T-CDCA, G-CDCA, T-DCA, G-DCA) in the liver, bile duct, gallbladder, upper intestine, lower intestine, colon, portal vein, intrahepatic sinusoidal space and systemic circulation, as well as the dynamics of FGF19 to model the inhibition of BA synthesis in the liver. Voronova *et al*. explicitly represent mealdependent gallbladder emptying and intestinal transit, ileal absorption, hepatic uptake, microbial deconjugation/dehydroxylation, and feedback regulation of BA synthesis through intestinal FXR–FGF19 and hepatic FXR signaling. These are particularly relevant because the model separates physiological parameters such as intestinal motility and blood flow from BA-specific parameters such as synthesis, absorption, and hepatic uptake. Hence, it provides an ideal foundation for patient-specific parameter inference.

### Computational barriers to patient-specific EHC modeling

Realizing the clinical potential of a personalized EHC model requires efficient estimation of the different parameters from clinically measured data. To do so, two main challenges arise. Firstly, the EHC, like many biological processes within the human body [11, 12], is multiscale, spanning a wide range of time and length scales [13]. A consequence of mechanistic ODE models that incorporate multiple timescales is numerical stiffness. Definitions of stiffness are vague and imprecise, but in general a system of ODEs can be described as stiff if it has *“a particular solution that is smooth and slowly varying (relative to the time interval of the computation), but in a context where the nearby solution curves are much more rapidly varying”* [14]. The stable integration of a stiff system forces explicit solvers to take prohibitively small time steps, making every simulation computationally expensive.

Secondly, biological ODE models heavily rely on parameters that must be estimated from experimental data. Traditional parameter optimization techniques often yield singular point estimates for the values of these parameters, which fail to capture the inherent uncertainty and inter-individual variability typically present in biological measurements. In contrast, Bayesian inference offers a robust probabilistic framework to overcome this by treating the model parameters as random variables rather than fixed values, quantifying both estimation uncertainty and inter-individual variability through posterior distributions [15, 16, 17]. However, these posteriors are typically explored by Markov chain Monte Carlo (MCMC) sampling [15, 16, 17], which requires thousands of model simulations for convergence. Combined with the long simulation time caused by stiffness, Bayesian inference of the full EHC model becomes computationally prohibitive.

### Aims of this study

Realizing the clinical potential of a personalized EHC model, however, requires not just computational feasibility, but also that the clinical measurements used for inference are sufficiently informative themselves. Current sampling protocols for BA quantification are limited by logistical convenience rather than identifiability considerations: a single fasting peripheral blood draw is the clinical standard, while portal blood sampling is restricted to collection during surgery, and multi-time point dynamic protocols remain rare due to their burden on both patients and clinicians. It remains unclear whether these standard, single time point protocols contain sufficient information to uniquely identify the underlying model parameters, or whether multi-time point or multi-compartment sampling is required to uncover the complex, interacting processes of the EHC. Committing to a protocol without first establishing its identifiability properties exposes the study to the collection of data that cannot answer the intended question whilst simultaneously contributing to unnecessary clinical risk and analytical expenses. By simulating *in silico* patients and systematically comparing the posterior information yielded by alternative sampling strategies, computational approaches can identify a minimally burdensome protocol that achieves robust parameter identifiability before any patient is sampled.

To overcome these computational barriers and enable the clinical application of a personalized EHC BA model, our study aims to (1) develop a generalizable computational pipeline, combining quasi-steady state approximation (QSSA) model reduction with NN-derived surrogates, that makes the Bayesian inference of parameters feasible in stiff, high-dimensional mechanistic models; (2) establish baseline parameter identifiability by applying this pipeline to the human EHC model, which will both determine which parameters governing BA homeostasis can be identified from a single fasting blood draw, and characterize the structural degeneracies that limit individual-level inference; and (3) identify the optimal sampling protocol by systematically comparing protocols in which we varied the number of peripheral time points and the inclusion of portal blood measurements to determine the minimal clinically feasible protocol that resolves these degeneracies and achieves robust multi-parameter identifiability.

## Materials and Methods

### Model Reproduction and Refinement

The human EHC model of Voronova *et al*. [10] was reproduced in Python (version 3.12.4). The EHC model describes the amounts of each BA species in each compartment of Fig. 1 as a system of ODEs (Eq. 1):

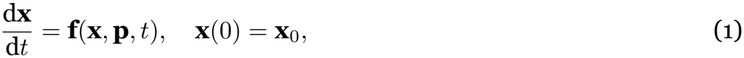

where **x**(*t*) *∈* ℝ^74^ is the vector of dynamical variables and **p** is the vector of model parameters. The state variables comprise the nine BA species (CA, CDCA, DCA, T-CA, G-CA, T-CDCA, G-CDCA, T-DCA, G-DCA) in the liver, bile duct, gallbladder, upper intestine, lower intestine, colon, portal vein, sinusoidal space, and systemic circulation, together with the variables of the FXR-FGF19-C4 feedback loop. The explicit time dependence in **f** arises from meal-dependent functions that drive gallbladder emptying and intestinal transit, which makes the system non-autonomous.

The parameters in vector **p** fall into four classes: (1) physiological quantities, such as compartment volumes and hemodynamic blood flow rates; (2) transport and kinetic rate constants, governing hepatic uptake, active ileal absorption, passive diffusion, and microbial transformations (deconjugation and 7*α*-dehydroxylation); (3) BA synthesis rates, specifically `ksyn_ca` and `ksyn_cdca`; and (4) Regulatory parameters, governing the FXR-FGF19 feedback on bile acid synthesis. Inter-individual variability in intestinal and colonic motility and in lower-intestinal absorption are represented by the scalars `intm_bsv`, `colm_bsv`, and `lint_abs_bsv` (Fig. 1). The complete set of ODEs and the parameters used can be found in S1 File and S2 File respectively. The Python code to reproduce the model (`voronova_model_reproduction.py`) can be found in a GitHub (GitHub, Inc., USA) repository (https://github.com/sebjoseph122/EBAP).

The regulatory feedback loop involving the inhibition of 7*α*-hydroxy-4-cholesten-3-one (C4) production, a precursor of CA and CDCA by FGF19, was modified by utilizing an inhibitory function as opposed to an exponential decay function, which better reflects receptor-mediated pharmacokinetics. The modification ensures C4 is non-negative and bounded (0 < *C*4 ≤*C*4_*max*_) and that the negative feedback mechanism is preserved since C4 synthesis always decreases if FGF19 amounts increase and vice versa (Eq. 2). The derivation of the equations and the parameters can be found in S3 File, while the modified model code can be found in `full_model.py`.

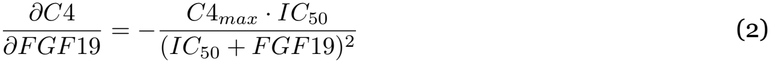

A simulation was conducted for both the original and modified model over a period of 1680 hours (70 days), starting at 08:00 on day 0 with meal intake simulated at 0.5, 4, 10 and 12.5 hours after 08:00 every day (08:30, 12:00, 18:00, 20:30) as performed by Voronova *et al*. [10]. A period of fasting was implemented after the last meal at 20:30 on day 68 (1644.5 hours). The initial condition of each variable was 0 and the solutions were recorded at 5041 time points (3 points per hour). The relative and absolute tolerances were kept at their default values: 1e-3 and 1e-6 respectively. The sum of the total BA pool across all compartments throughout the duration of the simulation was calculated for each model and compared as a means of validation for the new inhibitory feedback loop.

### Computational Performance Analysis

To evaluate the computational cost of solving the modified ODE model, the system was integrated using the `solve_ivp` function from the SciPy library [18]. To evaluate whether the model exhibits stiffness, it was solved using all the standard available integration methods, including explicit methods: the Runge-Kutta method of order 5(4) (`RK45`), the Runge-Kutta method of order 3(2) (`RK23`) and the Runge-Kutta method of order 8 (`DOP853`); implicit methods: the Runge-Kutta method of the Radau IIA family of order 5 (`Radau`) and the multi-step variable-order (1 to 5) method based on a backward differentiation formula for the derivative approximation (`BDF`); and the Adams/BDF method with automatic stiffness detection and switching (`LSODA`). In the context of numerically stiff systems, explicit methods are forced to take a large number of small time steps since the local error is dominated by what happens at the fast time scale. Taking larger time steps leads to instability and increases oscillations. Consequently, ODE systems have also been described as stiff if *“explicit methods don’t work”* [19, 20].

Simulations were conducted as described above (see Model Reproduction and Refinement). All simulations were performed on a local workstation (13th Gen Intel^®^ Core™ i7-1355U processor). The performance of each solver was evaluated on the total time required to complete the simulation, the number of times the solver evaluated the ODE system and the number of successful and failed time steps taken to reach the final time point in a Jupyter Notebook (`manuscript_notebook.ipynb`). For reliability, each integration method was performed in a new Python kernel in triplicate, and the average of the total time for each simulation was calculated.

### Stiffness Analysis

Mathematically, the stiffness ratio of an ODE system can give some understanding of whether a system is stiff [14]. This is the ratio between the magnitudes of the real parts of the largest and smallest eigenvalues over all *i* eigenvalues of the Jacobian matrix (Eq. 3).

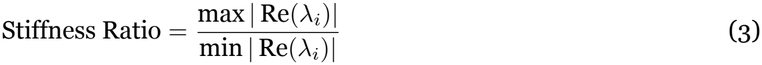

When this ratio spans several orders of magnitude, the system can be considered stiff; though there are some scenarios where this does not hold true. For example, additional time scales may come from inhomogeneous forcing terms and time-dependent coefficients, that are not captured by the eigenvalues, or a system may be stiff over some time intervals and not stiff elsewhere [14].

To quantify the degree of numerical stiffness throughout the 70-day simulation, the Jacobian matrix, **J**(*t*, **x**), was calculated numerically using a forward finite difference approximation (Eq. 4):

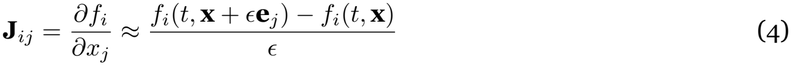

where **x** is the vector of state variables, *t* is the simulation time, *ϵ* is a small relative perturbation (1e-6), and **e**_*j*_ is the *j*-th unit vector.

Due to the time-dependent meal functions, stiffness was analyzed at five physiological time points: 0, 1, 22, 1609, and 1660 hours (`stiffness_analysis.ipynb`). These points enable the tracking of numerical stiffness across the initial condition (*t* = 0), the first meal transient (*t* = 1), a fasting period before reaching the regular limit cycle of the system (*t* = 22), peak digestion within the established limit cycle (*t* = 1609), and the final extended fasting period (*t* = 1660). The numerical stiffness at each point was characterized by the Stiffness Ratio (Eq. 2). The eigenvalues, *λ*_*i*_, of the Jacobian matrices at these time points were computed using the `numpy.linalg.eigvals` library. A high stiffness ratio indicates a significant separation of timescales. To identify the physiological drivers of this stiffness, participation factor analysis was performed on the corresponding eigenvectors as described by Garofalo *et al*. [21].

Participation factors (*p*_*ki*_) were calculated as the product of the *k*-th elements of the right (*u*_*i*_) and left (*v*_*i*_) eigenvectors corresponding to the *i*-th eigenvalue, such that 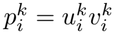. For each eigenvalue, the state variable demonstrating the largest participation degree was designated as the dominant variable for that timescale, identifying the primary candidate variables for the QSSA.

### Quasi-Steady State Approximation

One method to overcome the problem of numerical stiffness is to use the QSSA to reduce the ODE system by removing the ODEs representing the dynamics of fast variables, *x*_*i*_, by assuming they reach steady state instantaneously compared to the slow variables (Eq. 5), and then expressing them algebraically in terms of the slow variables [22, 23]. The resulting differential algebraic equation (DAE) system is a combination of ODEs for the slow variables and algebraic equations for the fast variables, *x*_*i*_, and has reduced computational overhead since the solver no longer has to take a large number of small time steps to account for the fast dynamics.

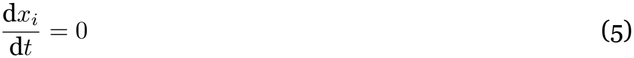

The ODEs representing the portal vein and sinusoidal BA species were approximated to be at steady state and set equal to zero. These ODEs were removed from the system and calculated algebraically. The derivation of the algebraic solution of these BA variables can be found in S4 File. The reduced model can be found in `qssa_model.py` and the code to calculate portal vein and sinusoidal BA during post-processing can be found in `manuscript_notebook.ipynb`.

To quantitatively validate the QSSA-reduced model, its trajectories were interpolated onto the full model’s time grid using cubic splines and compared state-by-state. Denoting the trajectory of state *i* in the full and reduced models by 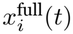 and 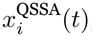, agreement was assessed by the local relative error (Eq. 6):

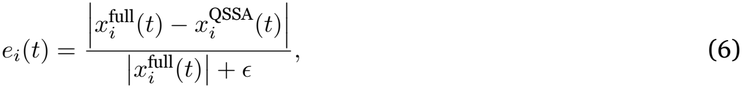

and by the integrated relative *L*_2_ norm (Eq. 7):

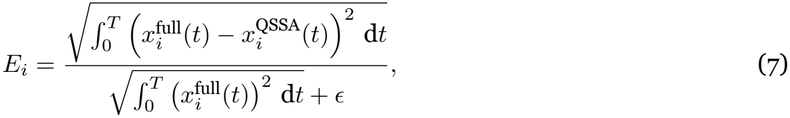

where *T* = 1657 h, *ϵ* = 10^−9^ is a regularization constant added to prevent numerical instability, and the integrals were evaluated using Simpson’s rule. Computational performance of the reduced model was then evaluated following the same protocol detailed in the Computational Performance Analysis section (`manuscript_notebook.ipynb`).

### Sensitivity Analysis

To identify which parameters impact fasting BA measurements the most and to guide the selection of parameters for high-dimensional inference, a local sensitivity analysis based on the Fisher information matrix (FIM) was performed [24]. The sensitivities were evaluated with respect to the fasting concentrations of the nine peripheral and nine portal BAs (*x*_*i*_, *i* = 1, …, 18), sampled at the end of a 12-hour fasting period (*t* = 1657 h), representing a single fasting blood draw rather than a dynamic postprandial time course. For each of the 75 continuous parameters *θ*_*j*_, the log-log sensitivity of each of the 18 BA concentrations was computed by finite-difference perturbation of the QSSA-reduced model (Eq. 8):

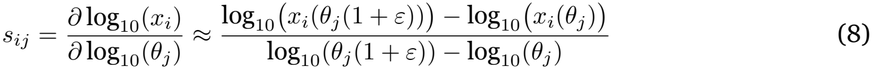

uated at the baseJline parameter values. Parameters were then ranked by the *ℒ*_2_ norm of their sensitivity where *ε* is a small relative perturbation and *x*_*i*_(*θ*_*j*_) denotes the fasting concentration of the *i*-th BA evalvector,evaluated at the baseline parameter values. Parameters were then ranked by the *ℒ*_2_ norm of their sensitivity vector,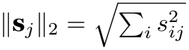, which quantifies the overall influence of each parameter on the 18 observable fasting concentrations (`fim.py`).

To assess whether the selected parameters could be jointly estimated without strong collinearity, the FIM was constructed as **F** = **S**^T^**S**, where **S** is the sensitivity matrix restricted to the selected parameters, and the expected pairwise correlation between parameters *j* and *k* was derived from the Cramér-Rao covariance matrix **C** = **F**^−1^ (Eq. 9):

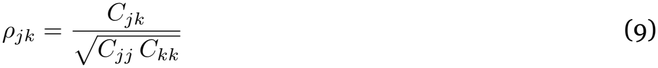

This correlation matrix was used to predict which parameters would exhibit posterior collinearity before running the full MCMC inference.

Five parameters were selected for high-dimensional inference based on both physiological relevance and sensitivity ranking. The selected parameters: the hepatic synthesis rates (`ksyn_ca`, `ksyn_cdca`) and the motility and absorption variability scalars (`intm_bsv`, `colm_bsv`, `lint_abs_bsv`) represent the core synthesis-motility-absorption axis of the EHC and all ranked within the top 13 most sensitive parameters out of 75. Parameters that ranked higher in sensitivity but were excluded corresponded to anatomical volumes, blood flow rates, and fixed kinetic parameters that are generally conserved across individuals in population-based modeling.

### Bayesian Inference Framework

Bayesian inference utilizes Bayes’ theorem, which states that the posterior probability distribution of the parameters (***θ***) given the experimental data (*D*), *P* (***θ***| *D*), is proportional to the product of the likelihood, *P* (*D*| ***θ***), and the prior probability, *P* (***θ***) (Eq. 10). The prior represents our initial belief or knowledge about the biological parameters before observing the data, while the likelihood quantifies how well the model explains the observed data.

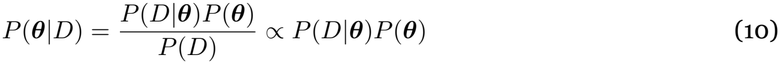

The resulting posterior distributions enable both the quantification of the uncertainty of parameter estimates and the identification of structural parameter identifiability. The log-likelihood function (ln ℒ) was evaluated assuming independent Gaussian noise in log-space (Eq. 11):

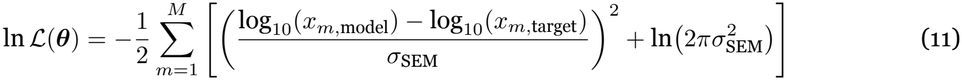

where *M* represents the total number of measured BA concentrations determined by the specific sampling protocol, *x*_*m*,model_ represents the *m*-th BA concentration predicted by the model or surrogate used, *x*_*m*,target_ is the *m*-th BA concentration observed in the synthetic data, and *σ*_SEM_ = 0.1 is the standard error of the mean in log-space.

### Neural Network-Derived Surrogate Modeling

Evaluating the likelihood function with the QSSA-reduced ODE model in a higher-dimensional space becomes computationally prohibitive due to the sheer number of evaluations required for MCMC convergence. To overcome this bottleneck, neural network surrogates were trained to map the parameter space directly to the model’s outputs [25], replacing the ODE system within the MCMC workflow and providing almost instantaneous likelihood evaluations [26].

To systematically compare sampling protocols, four separate multi-layer perceptron (MLP) surrogates were constructed, one for each sampling protocol (see Virtual Populations and Sampling Protocols below). For each surrogate, training data was generated using Latin Hypercube Sampling (LHS) to efficiently and uniformly map the five parameter (5P) space. A total of 12,000 parameter combinations were sampled in log_10_ space (`generate_lhs_5p.py`). The QSSA-reduced model was simulated for each combination via HPC array jobs to establish the deterministic mapping between the five log_10_ parameter inputs and the log_10_ target BA concentrations specific to a given sampling protocol.

Each MLP regressor was trained using the `scikit-learn` library. Both input features (log_10_ parameters) and output targets (log_10_ concentrations) were normalized using Z-score normalization prior to training. The optimized network architecture consisted of three hidden layers with 256, 256, and 128 nodes, utilizing a rectified linear unit (ReLU) activation function and the Adam solver.

Each surrogate was evaluated on two independent holdout sets generated with distinct random seeds: a 5P set in which 2,000 LHS samples across the full parameter space were drawn and a one parameter (1P) set in which 300 samples where only `ksyn_ca` varied were drawn. All surrogates reproduced the ODE outputs with high fidelity, yielding Root Mean Square Errors (RMSE) of *∼*0.010-0.012 in log_10_ space. Because these surrogate errors are an order of magnitude smaller than the assumed observational noise (*σ*_SEM_ = 0.1), the MLPs were deemed statistically valid substitutes for the QSSA-reduced ODE model within the `emcee` log-likelihood evaluation loop.

### Surrogate Validation via single parameter inference

Before deploying the surrogate for high-dimensional inference, it was critical to verify that the NN approximation does not bias the inferred posterior distributions. We validated the surrogate by replicating the single parameter Bayesian inference of `ksyn_ca` using both the QSSA-reduced ODE model and the NN surrogate. The single time point peripheral and portal sampling protocol (18 targets) was used as a test case, with the remaining four parameters fixed to their baseline values.

### Virtual Populations and Sampling Protocols

To evaluate parameter identifiability, *in silico* experiments were conducted using virtual populations. Two populations of 100 virtual patients were generated. For the 1P validation, only `ksyn_ca` was varied, sampled from a log-normal distribution (*µ*_log_ ≈1.44, *σ*_log_ = 0.15), while the other four parameters were fixed to their baseline values. For the 5P validation, all five selected parameters were sampled independently from log-normal distributions (*σ*_log_ = 0.15) centered at their baseline values.

To determine the minimal sampling protocol required for robust identifiability, four experimental protocols were evaluated (Table 1). For all protocols, synthetic observations were generated by simulating the QSSA-reduced model for each virtual patient and adding log-normal noise (*σ*_SEM_ = 0.1).

**Table 1:** Summary of the four experimental sampling protocols evaluated for parameter identifiability. Each protocol samples nine bile acid species (CA, CDCA, DCA, and their taurine- and glycine-conjugated forms) in the indicated compartments.

| **Design** | **Compartments sampled** | **Sampling times (h)** | **Targets (*M*)** |
|---|---|---|---|
| **Peri 1tp** | Peripheral | 1657 | 9 |
| **PeriPort 1tp** | Peripheral + Portal | 1657 | 18 |
| **Peri 3tp** | Peripheral | 1644.5, 1650.75, 1657 | 27 |
| **Peri 5tp** | Peripheral | 1644.5, 1647.625, 1650.75, 1653.875, 1657 | 45 |

### MCMC Inference and Evaluation Metrics

For each of the four experimental protocols, Bayesian inference was performed using both the 1P and 5P virtual populations, resulting in eight inferences. The 1P method tests whether the sampling protocol contains sufficient information to identify a single parameter in isolation, while the 5P method tests whether parameters can be inferred jointly and reveals structural degeneracies.

The posterior distributions were explored using a parallelized MCMC architecture utilizing the affine-invariant ensemble sampler implemented in the `emcee` package [27]. All inference was executed using a parallelized multiprocessing pool, with the 100 independent patient fits distributed across available CPU cores. The single parameter inference using the QSSA-reduced ODE model was performed on a high-performance computing (HPC) cluster (24 CPU cores, single node). All surrogate-based inference runs were executed on a local workstation. To ensure a fair and unbiased comparison across protocols, all 5P inference runs utilized identical, MCMC settings: 32 ensemble walkers were evolved for 6,000 steps, with a burn-in period of 1,500 steps discarded to ensure the chains reached a stationary distribution. Walker starting positions were drawn uniformly from the prior bounds to avoid initialization bias. Uninformed uniform priors were applied, matching the LHS training bounds.

Parameter recovery and identifiability were assessed using several metrics: (1) the coefficient of determination (*R*^2^) between the per-patient posterior means and the true virtual patient values for quantifying the fraction of inter-individual variability captured by the inference; (2) the Mean Absolute Error (MAE) and RMSE in log_10_ space for quantifying individual-level accuracy; (3) the 95% credible interval (CI) coverage, defined as the fraction of true parameter values falling within the 2.5th and 97.5th percentiles of the posterior distribution for uncertainty calibration; and (4) the median within-patient posterior correlation matrix, computed to quantify parameter degeneracy across the virtual population.

## Results

### The inhibitory Hill function feedback loop reproduces meal-response dynamics

Before the model could be used for stiffness analysis and parameter inference, we first verified that the modification to the FGF19 induced inhibitory feedback loop of BA synthesis did not alter the fundamental dynamics of the Voronova *et al*. model. The original model represents the inhibition of hepatic C4 production by FGF19 as an exponential decay. Instead, we replaced this with a bounded inhibitory Hill function, which ensures non-negative C4 and better reflects receptor-mediated pharmacokinetics (S3 File). The comparison between the original and modified models revealed that the modified model reaches a very similar limit cycle and a comparable fasting transient (Fig. 2A). Moreover, the predicted dynamics of BA concentrations during 24 hours of feeding followed by 24 hours of fasting across the compartments are qualitatively similar though their absolute values differ (Fig. 2B & Suppl. Fig. 1). Given these similarities, the refined model was considered to preserve the key dynamics and was therefore used for all subsequent analyzes and will be referred to as the full model from here.

**Figure 2.**
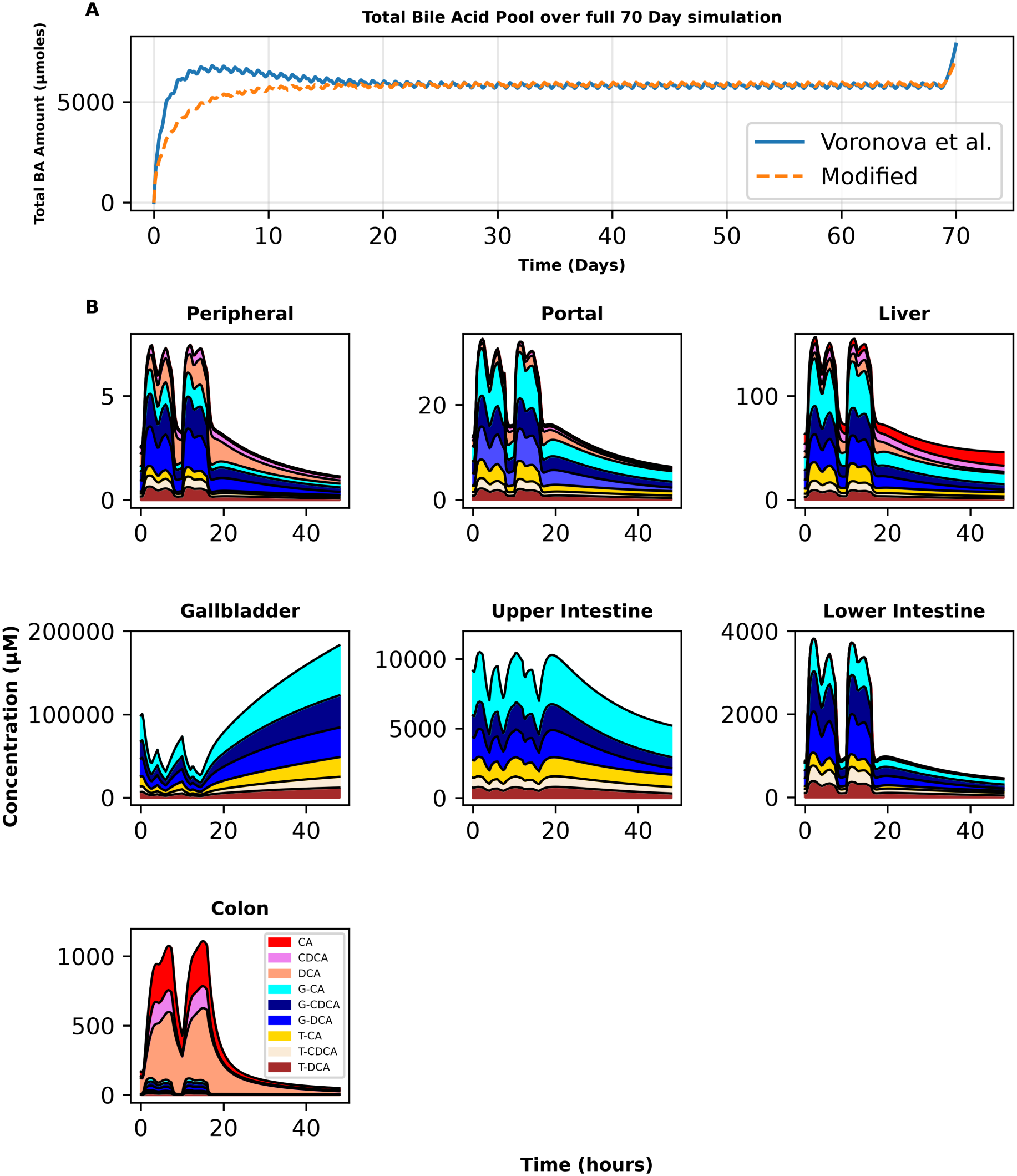
The ODE model of the enterohepatic circulation, modified with an inhibitory Hill function, captures the same limit cycle as the original model. (A) The total amounts of bile acids in all compartments of the original and modified models were summed respectively and plotted over time. (B) The predicted concentrations of 9 bile acids in the modified model were plotted including a 24 hour limit cycle followed by 24 hours of fasting in the peripheral blood, portal blood, liver, gallbladder, upper intestine, lower intestine and colon. Abbreviations: *CA, cholic acid; CDCA, chenodeoxycholic acid; DCA, deoxycholic acid; G-, glycine-conjugated; T-, taurine-conjugated*.

### The enterohepatic circulation ODE model exhibits numerical stiffness

To evaluate whether the full ODE model exhibits stiffness, we solved the model using the integrators available in `solve_ivp()`. While the full model can be solved using explicit methods RK45, RK23 and DOP853, the computational burden of using these methods is substantial, illustrated by the 5 to 8 times slower compute time compared to when solved with the LSODA integrator (Table 2), which automatically detects stiffness and switches between the Adams and BDF methods.

**Table 2:** Computational performance of various ODE solvers for a 70-day (1680 h) simulation of the enterohepatic circulation model with the modified feedback mechanism (Full Model). Solvers were evaluated based on execution time, function evaluations, and total successful steps.

| **Solver** | **Status** | **Time (s)** | **No. of evaluations** | **No. of Steps** |
|---|---|---|---|---|
| RK45 | **Success** | 429.82 | 8,253,890 | 1,178,616 |
| RK23 | **Success** | 298.99 | 4,670,327 | 1,553,322 |
| DOP853 | **Success** | 395.18 | 7,377,890 | 611,216 |
| Radau | Failed | 6.88 | N/A | N/A |
| BDF | Failed | 7.71 | N/A | N/A |
| LSODA | **Success** | **55.04** | **1,301,604** | **49,990** |

Furthermore, the disparity in computational performance is evident in the step count. While `RK45` and `RK23` require over 1.1 million steps for the 70 day simulation, `LSODA` completed the task in *∼*50,000 steps. This suggests that the explicit solvers are forced to use exceptionally small time steps to maintain stability during rapidly changing transients, a hallmark of a stiff system. Meanwhile, the dedicated stiff solvers `Radau` and `BDF` fail to converge, further highlighting the robustness of the `LSODA` algorithm for the 74-variable EHC ODE model.

The computational burden is further reflected in the total number of function evaluations. Explicit methods, such as `RK45`, exhibit a massive overhead, requiring over 8 million evaluations. This is six times the number required by `LSODA` (*∼*1.3 million), indicating that the explicit solvers frequently reject steps and requires infinitesimal step sizes to maintain numerical stability to account for the system’s fast variables.

Having established that the full model exhibits stiffness qualitatively, we evaluated the stiffness ratio to check whether the quantitative criteria for a system’s stiffness was met. The stiffness ratio was calculated across the simulation timeline. This revealed a massive separation of timescales, with the stiffness ratio consistently exceeding 4.6 ×10^4^ (Table 3). This confirms that the model meets the formal criteria for high numerical stiffness throughout all physiological phases.

**Table 3:**
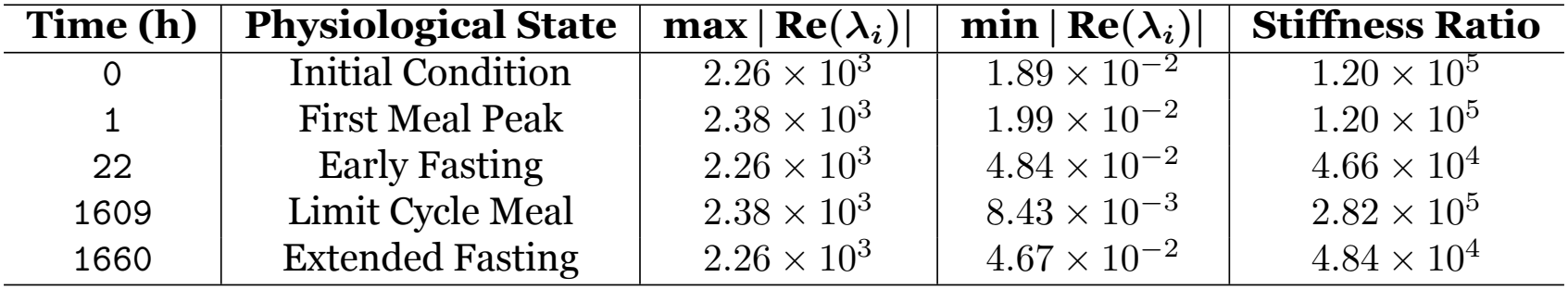
Eigenvalue magnitudes and resulting stiffness ratios at five key time points representing initial conditions, peak digestion, and basal fasting states.

| **Time (h)** | **Physiological State** | **max \| Re(λ<sub>i</sub>)\|** | **min \| Re(λ<sub>i</sub>)\|** | **Stiffness Ratio** |
|---|---|---|---|---|
| 0 | Initial Condition | 2.26 × 10<sup>3</sup> | 1.89 × 10<sup>−2</sup> | 1.20 × 10<sup>5</sup> |
| 1 | First Meal Peak | 2.38 × 10<sup>3</sup> | 1.99 × 10<sup>−2</sup> | 1.20 × 10<sup>5</sup> |
| 22 | Early Fasting | 2.26 × 10<sup>3</sup> | 4.84 × 10<sup>−2</sup> | 4.66 × 10<sup>4</sup> |
| 1609 | Limit Cycle Meal | 2.38 × 10<sup>3</sup> | 8.43 × 10<sup>−3</sup> | 2.82 × 10<sup>5</sup> |
| 1660 | Extended Fasting | 2.26 × 10<sup>3</sup> | 4.67 × 10<sup>−2</sup> | 4.84 × 10<sup>4</sup> |

It is worth noting that the stiffness ratio fluctuates with regard to the 24-hour feeding cycle. While the basal fasting state (*t* ∈22, 1660) maintains a ratio between 4.66× 10^4^ and 4.84 ×10^4^, the introduction of meals significantly exacerbates the numerical stiffness. At the peak of the first meal transient (t=1), the ratio rises to 1.20× 10^5^, and reaches its maximum of 2.82× 10^5^ during the rhythmic limit cycle meal peak at t=1609.

### Sinusoidal space and portal vein are the main drivers of stiffness

Having established that the stiffness ratio is consistent during fasting periods, we identified the main drivers of the observed stiffness by performing participation factor analysis on the Jacobian matrix at *t* = 1660. This revealed that the system’s stiffness is localized within specific physiological compartments. By calculating the participation factors (*p*_*ki*_) for each mode, a clear gap separating the dynamics of the model was identified (Fig. 3).

**Figure 3.**
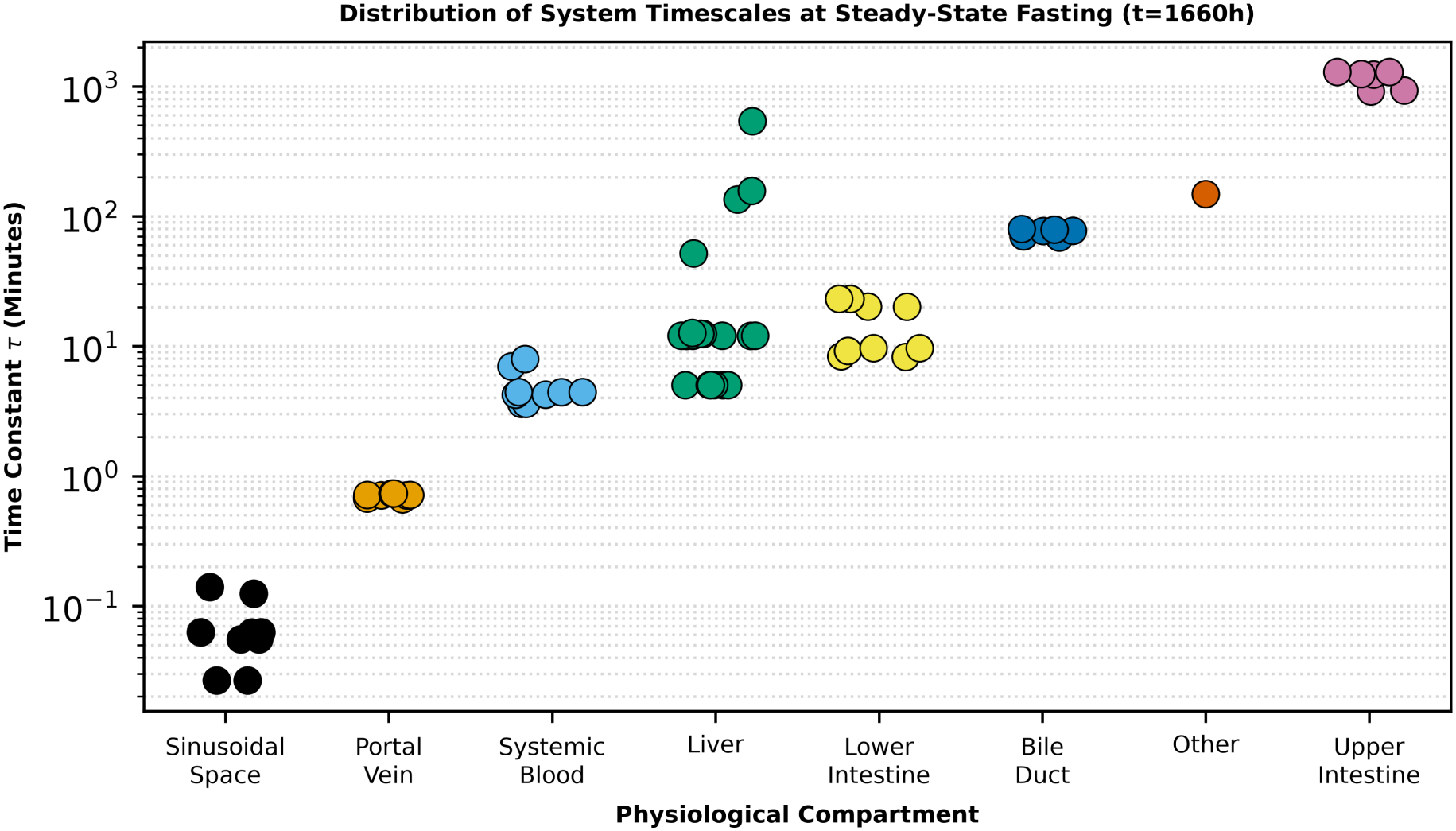
Distribution of characteristic system timescales across physiological compartments at fasting (t = 1,660h) after the establishment of a limit cycle. Each data point represents the time constant (*τ* = 1/ |Re(*λ*_*i*_) |) of an individual system mode. The dominant variable for each mode was identified using participation factors (*p*_*ki*_), which provide a dimensionless measure of state involvement in mode evolution. A significant spectral gap is observed between the fast transport dynamics of the portal vein and sinusoidal space (*τ*≈ 1.5–5 s) and the slower systemic and intestinal processes (*τ* ≈70–145 min), highlighting these small-volume compartments as primary candidates for the Quasi-Steady State Approximation (QSSA).

The stiffest variables in the system are those with time constants, *τ*, between 1.5 and 4 seconds which exhibit participation factors greater than 0.99 for BAs in the sinusoidal space and portal vein. This mathematical identification of sinusoidal and portal compartments as operating on a timescale of seconds is consistent with our knowledge of physiology. Physically, these eigenvalues represent the rapid transport of BAs within these small-volume compartments. Because these modes are almost exclusively driven by these specific state variables, they are the primary sources of the (4.84 ×10^4^) stiffness ratio that requires small integration steps when explicit solvers are used.

Conversely, BA dynamics in the liver and intestine are characterized by much longer time constants, ranging from 70 to over 140 minutes. This significant separation between these timescales, and those of the sinusoidal space and portal vein provides a mathematical justification for applying the QSSA. By setting the fast-participating variables in the sinusoidal space and portal vein to their algebraic equilibria, the ODEs representing the stiffest variables can be removed from the system while preserving the dynamics of the more slowly changing BAs in the liver and intestine.

### QSSA speeds up the compute time of the ODE system significantly

We used the QSSA to remove the ODEs representing the sinusoidal space and portal vein BAs by setting them equal to zero and solving for them algebraically (S4 File). The subsequent QSSA-reduced model was solved using the integrators available in `solve_ivp()`, revealing a significant reduction in the compute time by explicit solvers, with compute times of less than 10 seconds for `RK45`, `RK23` and `DOP853` (Table 4). This represents a percentage decrease of more than 97% for the three solvers, while the compute time for LSODA was halved from 55.04 seconds to 27.04 seconds. The reduction of compute time in these cases was a consequence of reducing model stiffness, which caused a significant reduction in the number of function evaluations and steps taken by the solver. Meanwhile, both the full and QSSA-reduced models failed to converge when the `Radau` and `BDF` solvers were used (Tables 2 and 4) because their finite-difference Jacobian estimates overflowed during the initial stiff transient, causing the BA concentrations in the stiff compartments to diverge to non-finite values.

**Table 4:**
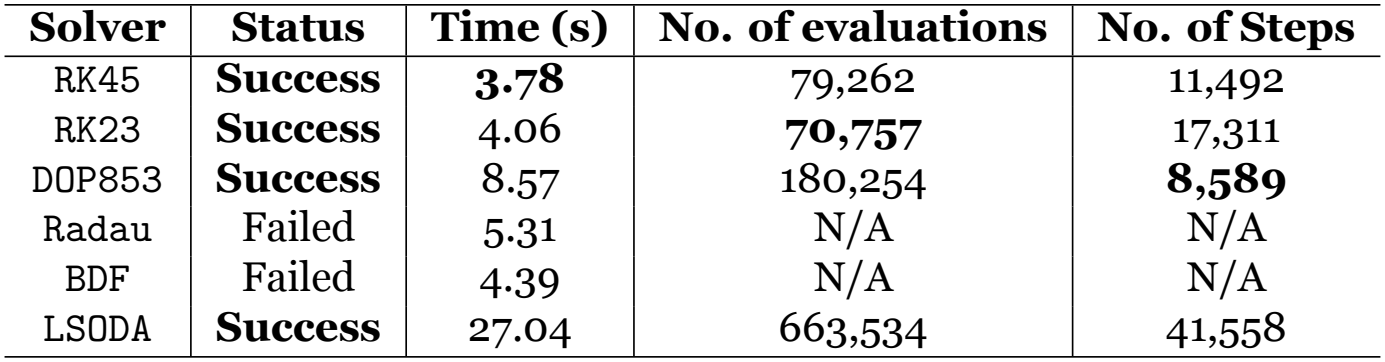
Computational performance of various ODE solvers for a 70-day (1680 h) simulation of the Quasi-Steady State Approximation reduced model of the enterohepatic circulation. Solvers were evaluated based on the execution time, function evaluations, and total successful steps. The time reported is the average of three simulation executions.

An analysis of the results of the simulation of the QSSA-reduced model revealed that the algebraic reconstruction of sinusoidal G-CA concentrations closely matches with the predictions from the full model during both the initial 72 hours (Fig. 4A) and the limit cycle (Fig. 4B). Furthermore, phase space projection confirmed that the QSSA-reduced model accurately traces the 24 hour limit cycle on the slow manifold, maintaining the relationship between a fast variable, G-CA_*SIN*_, and a slower variable CA_*COL*_ (Fig. 4C). Evaluating how the error between the full model and the QSSA-reduced model across all BAs and across the 1,680 hour simulation demonstrated long-term mathematical stability (Fig. 4D). The error initially spikes from 0 to 17.5%, but this quickly decays as both models reach a limit cycle, fluctuating between 0 and a meal-induced 5%. Calculating the integrated relative norm (ℒ _2_) results in 0.9029%, illustrating the high resemblance between the QSSA-reduced model and the full model.

**Figure 4.**
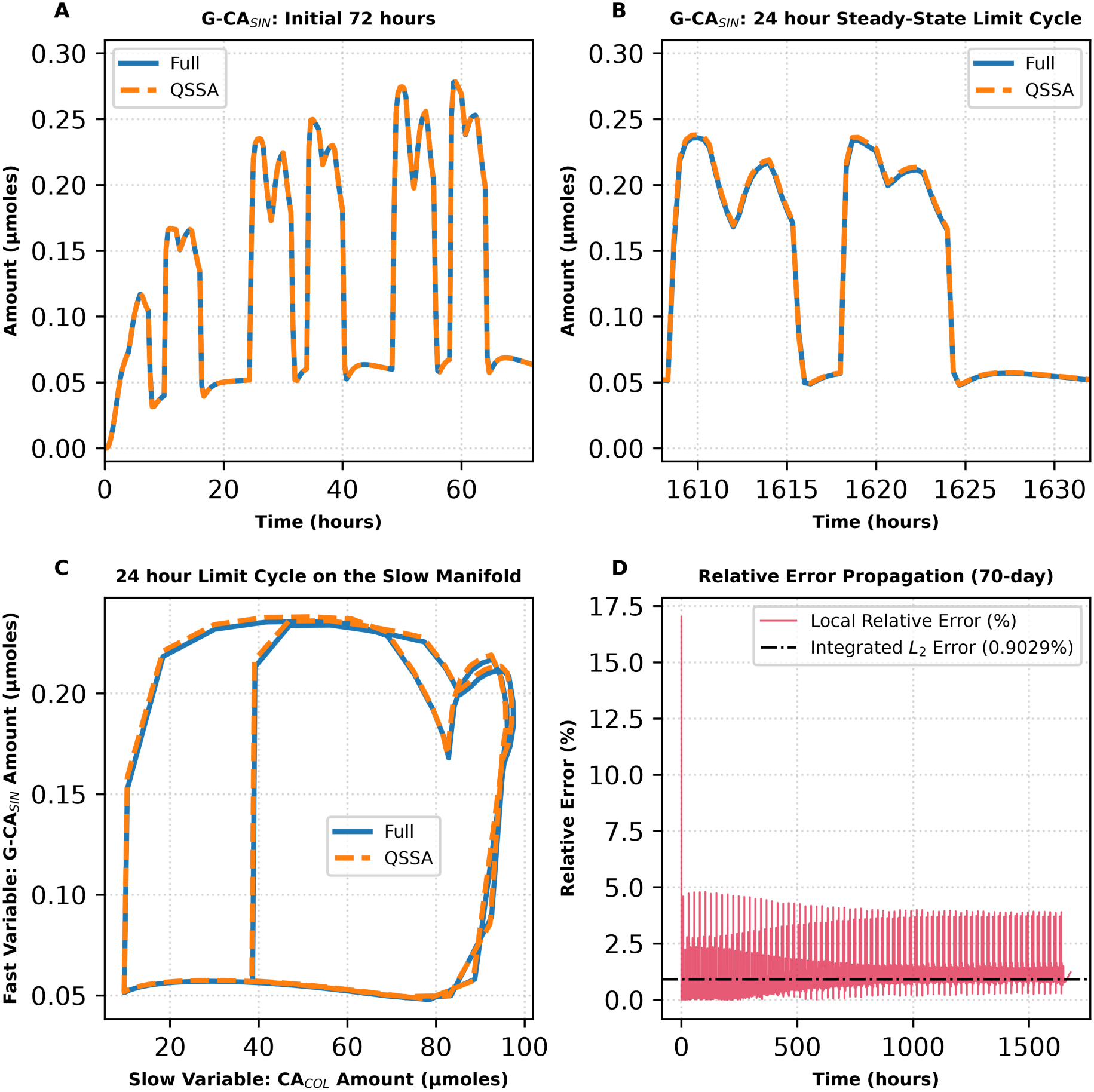
The quasi-steady state approximation reduced model retains similar dynamics to the full model with the inhibitory Hill function. (A) The trajectories of sinusoidal G-CA, a fast variable, over the first 72 hours were plotted for both the full model and the QSSA-reduced model. (B) The trajectories of sinusoidal G-CA during a 24 hour limit cycle were plotted for both models. (C) The 24 hour limit cycle on a slow manifold was visualized by plotting a fast variable, sinusoidal G-CA against a slow variable, colonic CA. (D) The local relative error between both models was evaluated over the course of a 1,680 hour simulation and plotted. Integration of this local error yields a global ℒ_2_ relative error norm of 0.9029%. Abbreviations: *CA, cholic acid; COL, colon; G-CA, glycocholic acid; QSSA, quasi-steady state approximation; SIN; sinusoidal space*.

### Bayesian inference recovers the population distribution of a single parameter

Since the QSSA-reduced ODE model reproduces the full model faithfully whilst substantially reducing the computational cost, we next assessed whether Bayesian inference could recover the known distribution of a single parameter from the fasting portal and peripheral BA concentrations at a single time point of a virtual population (*N* = 100). `ksyn_ca` was chosen for this initial one-dimensional validation because hepatic synthesis rates are a crucial determinant of BA homeostasis, are expected to vary between individuals, and are ranked within the most sensitive parameters in the FIM (Fisher information matrix) screening (Fig. 5), confirming that the fasting BA observations are informative about this parameter. The virtual population was generated by sampling `ksyn_ca`, from a log-normal distribution (*µ*_log_ = 1.44, *σ*_log_ = 0.15), and the fasting nine peripheral and nine portal BA concentrations were collected at *t* = 1657 h and log-normal noise (*σ*_SEM_ = 0.1) was added.

**Figure 5.**
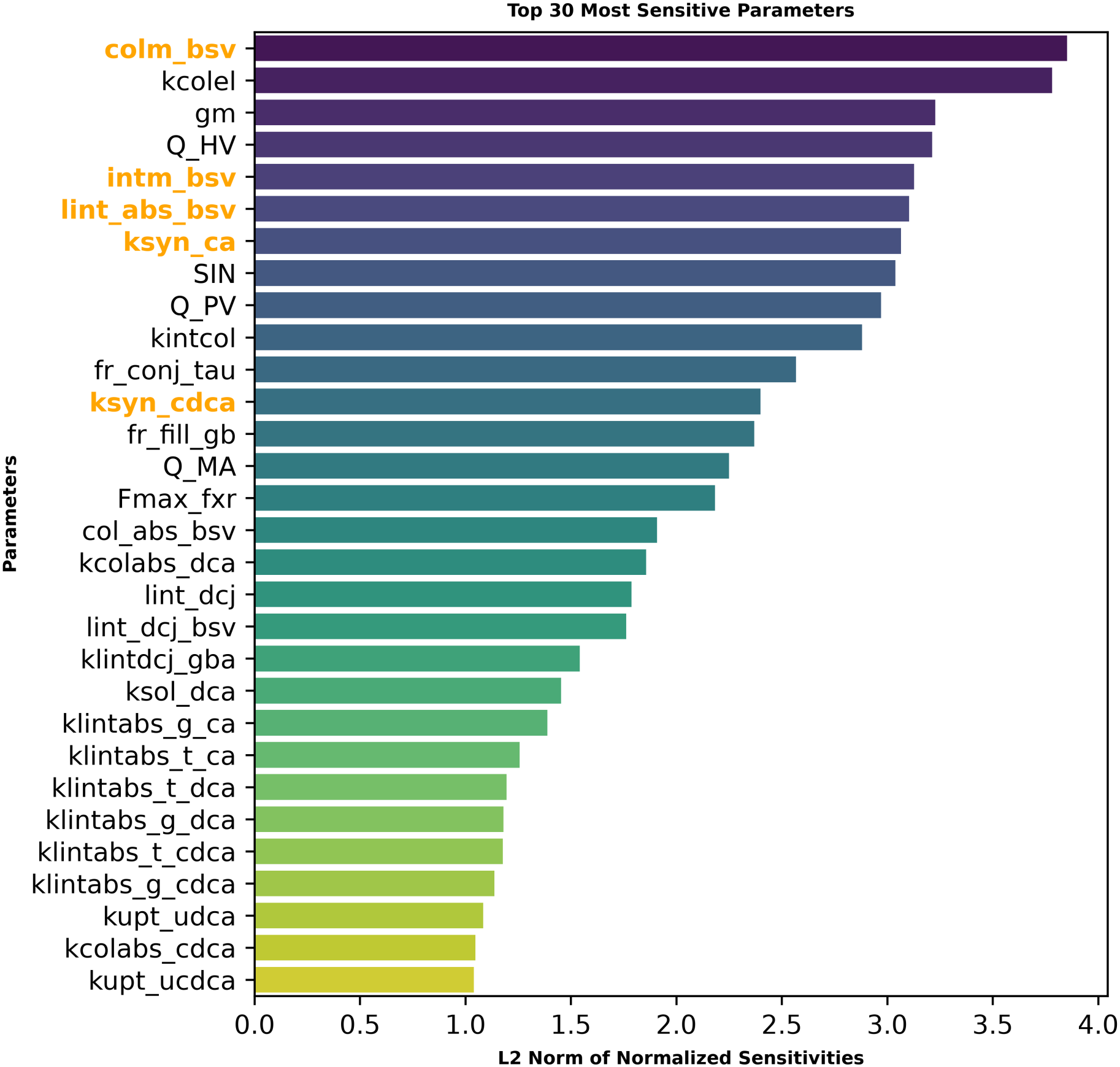
Fisher Information Matrix sensitivity analysis identifies the most influential parameters. Log-log sensitivities of the 18 fasting bile acid concentrations (9 peripheral and 9 portal) to each of the 75 model parameters, evaluated at *t* = 1657 h. Parameters are ranked by the ℒ_2_ norm of their sensitivity vector. The five parameters highlighted in orange (`ksyn_ca`, `ksyn_cdca`, `intm_bsv`, `colm_bsv`, `lint_abs_bsv`) all rank within the top 13 most sensitive parameters and were selected for joint Bayesian inference on the basis of both sensitivity ranking and physiological relevance to inter-individual variability. Parameters with higher sensitivity scores correspond to anatomical volumes, blood flow rates, and conserved kinetic constants that are conventionally held at population values. Descriptions of the parameters can be found in S2 File.

For each virtual patient, the posterior distribution of log_10_(`ksyn_ca`) was then sampled by MCMC with the QSSA-reduced model as the likelihood evaluator, using an uninformed uniform prior. The pooled posterior closely reproduced the true distribution, with *µ* = 1.420 and *σ* = 0.140 against the true values of 1.424 and 0.136 respectively (Fig. 6A).

**Figure 6.**
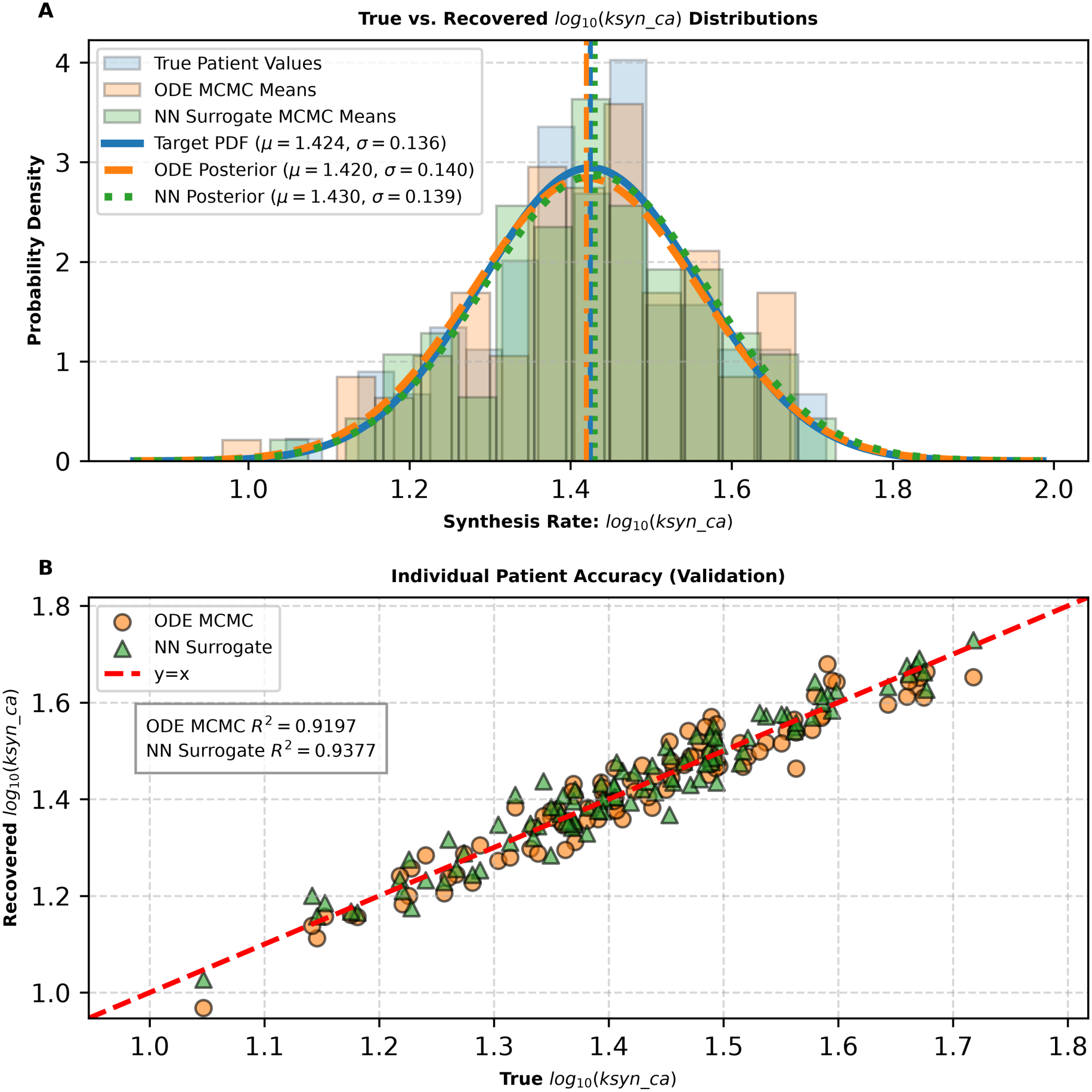
The neural network derived surrogate model reproduces the one parameter Bayesian inference performed with the QSSA-reduced ODE model. (A) Histograms of the true log_10_(`ksyn_ca`) values of the *N* = 100 virtual patients and of the posterior means recovered by MCMC inference with the QSSA-reduced ODE model and with the NN derived surrogate model were plotted, with the corresponding fitted normal probability density functions overlaid. Both inferences recovered the target distribution (*µ*_log_ = 1.44, *σ*_log_ = 0.15). (B) The posterior means of each patient’s log_10_(`ksyn_ca`) recovered by both models were plotted against the true virtual patient values, yielding *R*^2^ = 0.9197 for the QSSA-reduced ODE model and *R*^2^ = 0.9377 for the NN derived surrogate model. Abbreviations: *CA, cholic acid; ksyn_ca, hepatic synthesis rate of cholic acid; MCMC, Markov chain Monte Carlo; NN, neural network; ODE, ordinary differential equation; QSSA, quasi-steady state approximation*.

Moreover, parameter recovery at the individual patient level was very accurate, with the posterior means of each individual patient falling on or very close to the line of equality, with *R*^2^ = 0.9197 (Fig. 6B). Therefore fasting peripheral and portal BA concentrations at one time point were deemed to contain sufficient information to identify `ksyn_ca` at both the population level and the individual patient level. Having established this ODE-based ground truth, we next sought to accelerate inference by replacing the ODE solver with a neural network surrogate.

### A neural network surrogate reproduces the ODE-based inference

To eliminate the computational bottleneck posed by iterative ODE integration within the MCMC loop, we trained a multi-layer perceptron to map the 5P space directly to the log_10_ single time point fasting peripheral and portal BA concentrations. On an independent hold-out set of 2,000 Latin hypercube samples, the surrogate reproduced the QSSA-reduced model predictions with a RMSE of 0.010 log_10_ units and a maximum absolute error of 0.078, both below the assumed measurement noise (*σ*_SEM_ = 0.1). Accuracy was retained on a dedicated 1P validation set in which only `ksyn_ca` was varied (RMSE 0.010; maximum error 0.040), confirming that the surrogate interpolates accurately onto parameter subspaces not explicitly sampled during training.

Substituting the surrogate for the ODE system in the 1P inference reproduced the ODE-based results, yielding a posterior with *µ*≈ 1.430 and *σ*≈ 0.140 and an individual-level *R*^2^ = 0.9377 (Fig. 6), while reducing the cost of each likelihood evaluation by several orders of magnitude. Because the surrogate error is an order of magnitude smaller than the observational noise, it does not artificially tighten the posterior or introduce structural bias. The NN-derived surrogate was consequently adopted as the likelihood evaluator for all subsequent higher dimensional inference.

### Selection of five physiological parameters for inference

Next, we sought to expand the inference from a single parameter to a physiologically meaningful subset of the model’s 75 parameters. Sensitivity screening based on the FIM identified the parameters to which fasting peripheral and portal BA concentrations are most sensitive (Fig. 5). Five parameters were selected for joint inference on the basis of both their sensitivity and their physiological relevance to inter-individual variability: the hepatic synthesis rates of CA and CDCA (`ksyn_ca`, `ksyn_cdca`), the lower-intestinal absorption scalar (`lint_abs_bsv`), and the intestinal and colonic motility scalars (`intm_bsv`, `colm_bsv`). These parameters represent the inter-individual variability in the core synthesis-absorption-excretion pathway of the EHC, and all five ranked within the top 13 most sensitive parameters.

Although several parameters achieved higher sensitivity scores, the majority correspond to anatomical volumes, blood-flow rates and conserved fractional quantities that are conventionally held at population values, while the remainder are kinetic or regulatory rate constants that, in this model, do not encode inter-individual variability. Evaluating the likelihood with the QSSA-reduced ODE model across this 5P space was nonetheless computationally prohibitive because of the large number of model evaluations required for MCMC convergence.

### Multi-time point sampling resolves practical non-identifiability

To determine the minimal sampling protocol required for robust multi-parameter identifiability, we systematically compared four experimental protocols (Table 1): peripheral-only sampling at a single fasting time point (Peri 1tp, *M* = 9 targets), combined peripheral and portal sampling at a single fasting time point (Periport 1tp, *M* = 18), peripheral sampling at three time points during the extended fasting period (Peri 3tp, *M* = 27), and peripheral sampling at five time points during the extended fasting period (Peri 5tp, *M* = 45). For each protocol, parameter recovery was evaluated in a 5P virtual population (*N* = 100) using the surrogate-accelerated pipeline with identical, unbiased MCMC settings.

### 1P validation confirms practical degeneracies

Before evaluating joint identifiability, we confirmed that each protocol contains sufficient information to identify `ksyn_ca` in isolation by performing 1P inference with the other four parameters fixed at their baseline values. All four protocols recovered the true distribution of `ksyn_ca` with *R*^2^ > 0.88 (Suppl. Figure 2), confirming that the observed non-identifiability in the full 5P inference is not attributable to a lack of information in the data, but rather to practical degeneracies between simultaneously inferred parameters that arise because single time point measurements constrain only the net balance between synthesis and transit, not each process independently.

### Single time point experimental sampling protocols fail to resolve dynamic parameters

In the 5P inference, single time point sampling protocols exhibited severe practical non-identifiability for the dynamic parameters (Table 5; Suppl Fig. 3-4). For the Peri 1tp protocol, the coefficients of determination were *R*^2^ = 0.391 for `ksyn_ca`, *R*^2^ = 0.211 for `ksyn_cdca`, and *R*^2^ = −0.385 for `intm_bsv`, indicating that the recovered posterior means for intestinal transit were worse than simply predicting the population mean. The Periport 1tp protocol, which augments peripheral measurements with portal vein concentrations, slightly improved the coefficient of determination of `ksyn_ca` to *R*^2^ = 0.564 and improved recovery of `lint_abs_bsv` to *R*^2^ = 0.781 for, but failed to resolve intestinal transit (*R*^2^ = −0.135).

**Table 5:**
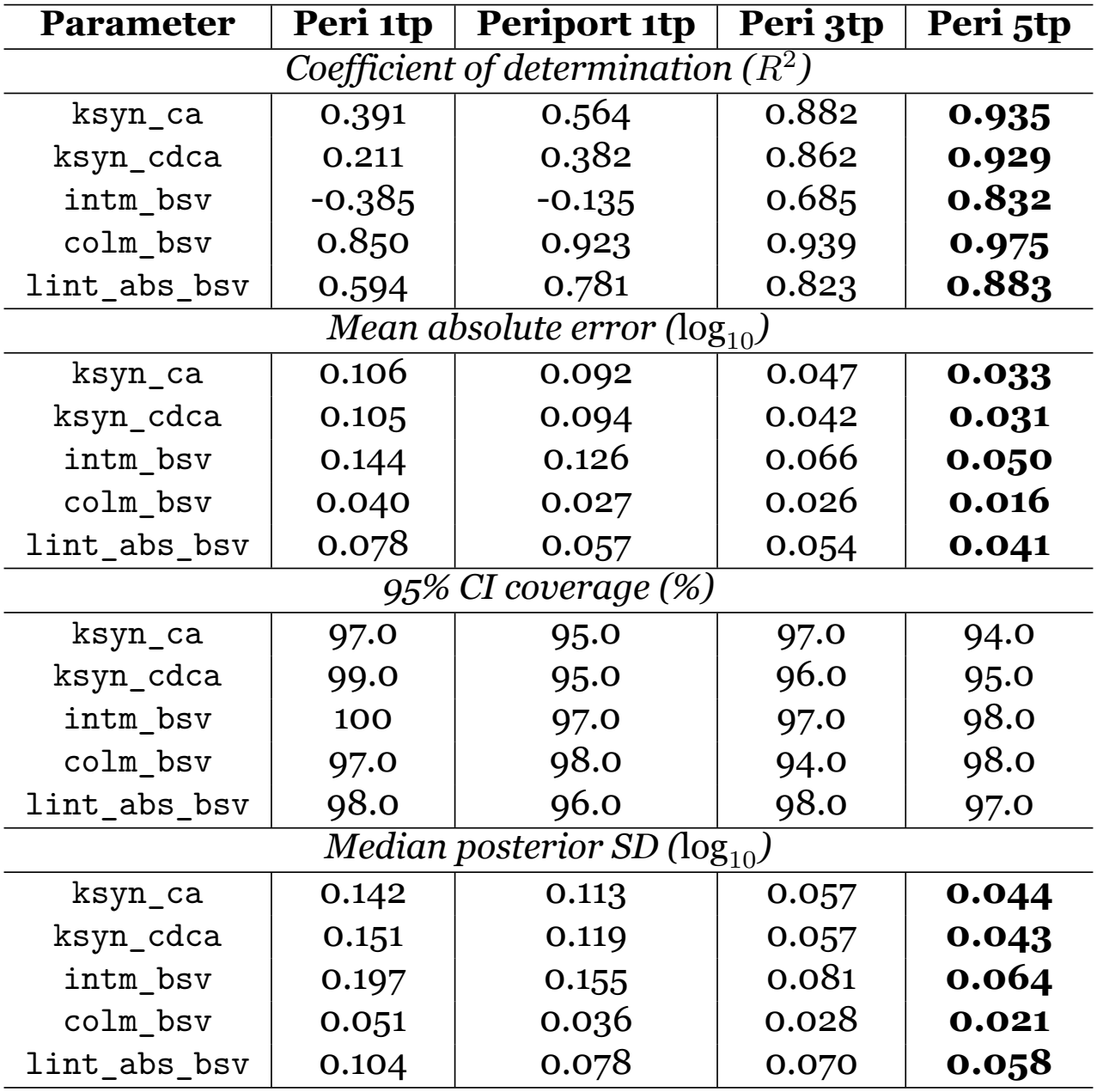
Five parameter recovery across the four experimental protocols. *R*^2^ and MAE are computed across the *N* = 100 virtual patients. Coverage indicates the fraction of true values falling within the 95% credible intervals. Posterior SD is the median across patients.

Crucially, the addition of nine portal BA concentrations did not break the synthesis-transit degeneracy: the median correlation between `ksyn_ca` and `intm_bsv` remained high with *r* = 0.83 compared to *r* = 0.79 for Peri 1tp protocol (Fig. 7), and the *R*^2^ for `intm_bsv` remained negative. This demonstrates that sampling an additional compartment at a single time point does not provide the information needed to disentangle synthesis from transit, despite the clinical invasiveness of portal blood collection.

**Figure 7.**
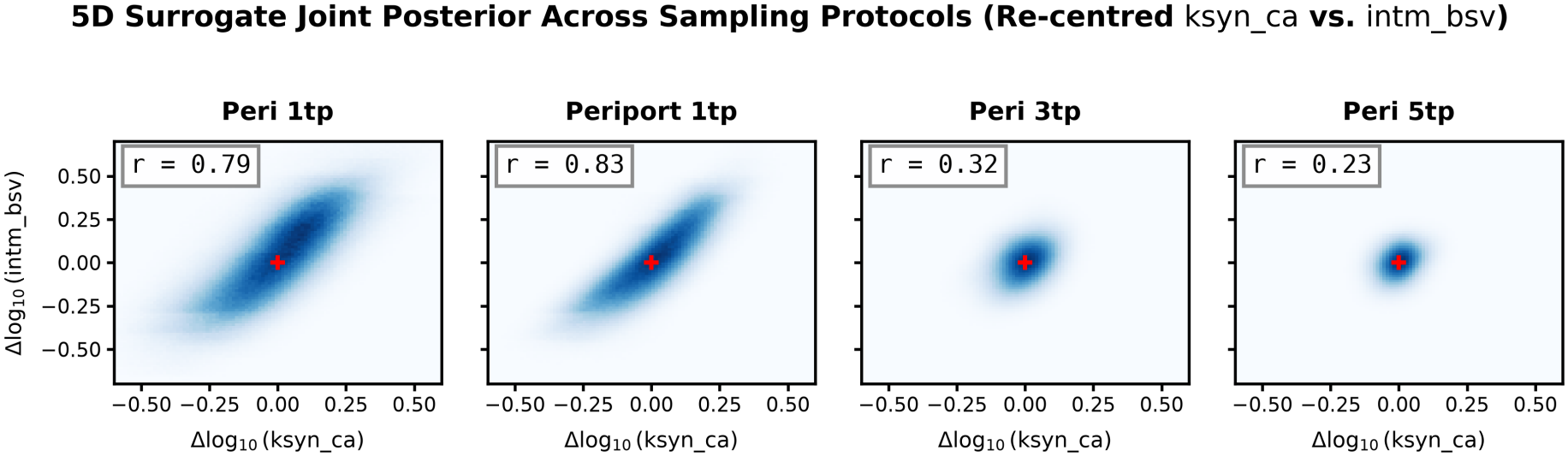
Temporal sampling breaks the synthesis-transit degeneracy. Median within-patient posterior correlation matrices for the four experimental protocols. The strong positive correlation between `intm_bsv` and `ksyn_ca` observed in single time point protocols (*r*≈ 0.8) progressively collapses with increasing temporal information (*r* = 0.33 at 3tp; *r* = 0.22 at 5tp), demonstrating that multi-time point sampling provides the orthogonal information needed to disentangle intestinal transit from hepatic synthesis. Abbreviations: *Peri, peripheral; Periport, peripheral + portal; tp, time point*.

The joint posteriors for both single time point protocols exhibited strong positive correlations between intestinal transit and hepatic synthesis with median *r* = 0.79 for Peri 1tp and *r* = 0.83 for Periport 1tp (Suppl. Figs. 3-4), suggesting that fasting BA concentrations at a single time point constrain only the net balance between synthesis and transit, not each process independently and hence confirming that the non-identifiability arises from a structural degeneracy. In addition, `intm_bsv` was correlated with lower-intestinal absorption `lint_abs_bsv` with median *r* ≈ 0.70 in both cases. In contrast, `colm_bsv` was only weakly coupled to the other parameters with median *r* ≤ 0.55, consistent with its identification across all four protocols (*R*^2^ ≥ 0.85).

### Multi-time point sampling breaks the synthesis-transit degeneracy

Multi-time point peripheral sampling resolved this degeneracy. The Peri 3tp protocol recovered all five parameters with *R*^2^ ≥ 0.68, including intestinal transit (Table 5), and reduced the median `ksyn_ca`-`intm_bsv` correlation from 0.83 to 0.32 (Fig. 7). The Peri 5tp protocol further improved recovery across all parameters (*R*^2^ ≥ 0.83), with the median `intm_bsv`-`ksyn_ca` correlation decreasing to 0.23 (Fig. 7). The progressive collapse of this correlation with an increasing number of time points demonstrates that observing temporal concentrations provides the information needed to disentangle synthesis from transit.

Importantly, the 95% credible interval coverage remained between 94% and 100% across all pro-tocols and parameters, indicating well-calibrated posteriors regardless of sampling protocol. The median posterior standard deviations decreased monotonically with increasing time point information: for `intm_bsv`, the median posterior SD declined from 0.197 log_10_ units (Peri 1tp) to 0.064 log_10_ units (Peri 5tp), reflecting the progressive resolution of the synthesis-transit degeneracy (Table 5).

Notably, while the primary synthesis-transit degeneracy was resolved, the posterior correlation between `intm_bsv` and `lint_abs_bsv` remained strongly positive. This positive correlation is a direct consequence of the compensatory nature of the EHC: to produce the same observed peripheral BA profile, a system with faster intestinal transit must compensate with a higher absorption efficiency (higher `lint_abs_bsv`) to prevent excessive fecal excretion. Conversely, slower transit provides greater contact time, allowing adequate net absorption even with a lower absorption rate scalar. Thus, this persistent correlation does not indicate a failure of the inference, but rather reflects the physiologically intuitive, compensatory parameter combinations required to match the observed data.

### Three time points represent a clinically efficient protocol

The Peri 3tp protocol recovered 92-96% of the *R*^2^ achieved by the Peri 5tp protocol for four of the five parameters, while reducing the number of required blood draws. Only `intm_bsv` exhibited a notable benefit from the additional two time points, recovering 82% of the Peri 5tp *R*^2^ at three time points compared to 100% at five. This is consistent with the identity of `intm_bsv` as a timing parameter that is most sensitive to the temporal resolution of the sampled trajectory. These results identify three peripheral time points as a clinically efficient compromise for multi-parameter BA phenotyping, while indicating that studies specifically targeting intestinal transit kinetics require the full five time point protocol.

## Discussion

BAs are tightly regulated metabolic signaling molecules whose enterohepatic circulation (EHC) is disrupted in several liver diseases. Our motivation is to understand how interacting physiological processes shape BA circulation and how alterations in these processes may account for the profiles observed in disease. As a step towards this goal, we investigate which physiological parameters can be recovered from different blood sampling protocols. Such mechanistic understanding could ultimately inform the evaluation and interpretation of circulating BA profiles as biomarkers of liver injury. However, the clinical utility of BAs remains limited by marked differences between portal and peripheral BA profiles and by substantial inter-individual variability arising from differences in EHC physiology. Mechanistic models are a highly valuable tool to infer inaccessible BA profiles and patient-specific metabolic parameters. Yet, personalization of existing EHC models remains computationally challenging because their multiscale dynamics cause numerical stiffness, while Bayesian parameter inference requires thousands of model simulations. To overcome these barriers, we developed a computational pipeline for Bayesian inference in a stiff, high-dimensional physiologically based model of the human EHC of BAs.

### Overcoming computational barriers to Bayesian inference

A major challenge in personalizing large mechanistic models is the computational cost of repeatedly solving stiff ODE systems during parameter inference. Here, we addressed this challenge through two complementary approaches: QSSA to reduce model stiffness and a NN surrogate to accelerate model evaluation. Together, these approaches reduced the computational cost of likelihood evaluation by several orders of magnitude, making ensemble MCMC inference across a virtual population of 100 individuals feasible within hours rather than weeks.

### QSSA reduces numerical stiffness while preserving EHC dynamics

Participation factor analysis identified the rapid dynamics of the portal vein and sinusoidal compartments as the main physiological sources of stiffness in the EHC model. Applying the QSSA to these fast variables removed the stiffest modes while preserving the slower hepatic and intestinal dynamics, with a relative L2 error of only 0.90% (Fig. 4). This reduced explicit-solver runtimes by more than 97% and approximately halved the runtime of LSODA (Table 4). Importantly, the fast variables were identified from a clear separation of physiological timescales rather than selected empirically, providing a mechanistic basis for the model reduction. These results demonstrate that timescale-based model reduction is a robust approach to making large, stiff pharmacokinetic models compatible with the large number of simulations required by Bayesian inference and other optimization techniques such as gradient-based optimization and genetic algorithms [28], whilst retaining their relevant biological dynamics.

### NN surrogate models enable high-dimensional Bayesian inference

Although QSSA substantially accelerated individual simulations, direct integration of the reduced ODE model remained too computationally expensive for high-dimensional MCMC inference. We therefore trained a NN surrogate to reproduce the mapping between the five selected parameters and the 18 fasting peripheral and portal BA concentrations. The surrogate reproduced the QSSA-reduced model with an RMSE of 0.010 log_10_ units, substantially below the assumed observational noise, and reproduced the one-dimensional `ksyn_ca` posterior obtained with the ODE model (Fig. 6). Thus, replacing ODE integration with the surrogate enabled extension from 1P to 5P inference at the population level.

The use of surrogate models nevertheless requires careful validation because their predictions are reliable only within the parameter domain represented during training. Here, agreement was evaluated using an independent global holdout set, a dedicated 1P validation set, and direct comparison of surrogate- and ODE-derived posterior distributions. This provides a general validation strategy for in-corporating NN surrogates into Bayesian inference of computationally intensive mechanistic models.

### Fasting BA concentrations reveal practical non-identifiability

The resulting pipeline accurately recovered the population distributions of all five parameters, with negligible bias and 95–98% credible-interval coverage (Table 5). However, parameter precision differed markedly at the individual level. Hepatic synthesis rates and intestinal motility showed broad posterior distributions and strong correlations (r = 0.83–0.90), while intestinal motility was also correlated with lower-intestinal absorption (r ≈ 0.70). In contrast, colonic motility was more tightly constrained.

These correlations reflect the homeostatic and compensatory nature of the EHC rather than a failure of the inference method. Fasting circulating BA concentrations result from the balance between synthesis, intestinal transit, absorption, and excretion. Consequently, changes in one process can be compensated by changes in another and produce similar circulating BA profiles. For example, reduced absorption may be compensated by increased hepatic synthesis, making these physiological states difficult to distinguish from fasting BA concentrations alone. The strong but incomplete parameter correlations therefore indicate practical non-identifiability: the available measurements constrain combinations of physiological processes more strongly than the individual parameters themselves.

These findings have direct implications for clinical study design. The practical non-identifiability revealed by single fasting measurements demonstrates that the standard single peripheral blood draw cannot distinguish between physiologically distinct states that produce similar circulating BA profiles. This limitation is not a failure of the inference method but a consequence of the homeostatic nature of the EHC: compensatory changes in synthesis, transit, and absorption can produce the same fasting peripheral BA profiles. Resolving this degeneracy requires measurements that capture the temporal dynamics of the system rather than a single equilibrium state.

### Dynamic sampling protocols resolve the synthesis-transit degeneracy

To determine whether temporal information can resolve the degeneracies identified above, we systematically compared four sampling protocols (Table 1), varying both the number of peripheral time points and the inclusion of a portal vein measurement. This comparison provides a quantitative roadmap for clinical study design. The negative coefficients of determination observed for `intm_bsv` under both single time point protocols (*R*^2^ = −0.385 for Peri 1tp; *R*^2^ = −0.135 for Periport 1tp) reflect the fact that the data contain no information about a parameter, the posterior collapses toward its prior mean and the recovered values are worse than simply predicting the population average. The strong positive correlations between `ksyn_ca` and `intm_bsv` (*r*≈ 0.8) in the single time point posteriors (Fig. 7) confirm that this degeneracy is structural rather than technical: fasting BA concentrations constrain only the net balance between hepatic synthesis and intestinal transit, not each process independently.

Adding portal vein measurements (Periport 1tp) modestly improved recovery of `ksyn_ca` (*R*^2^ = 0.564) and of `lint_abs_bsv` (*R*^2^ = 0.781), but failed to resolve `intm_bsv` ((*R*^2^ = −0.135). This finding has important clinical implications: portal blood sampling in humans is restricted to collection during surgery or invasive procedures such as the transjugular intrahepatic portosystemic shunt (TIPS) procedure, yet our results demonstrate that even this invasive approach, while being informative about hepatic BA exposure, does not yield sufficient information to disentangle synthesis from intestinal motility. These results argue against the clinical utility of portal sampling for multi-parameter inference.

In contrast, multi-time point peripheral sampling progressively broke the degeneracy. The Peri 3tp protocol recovered all five parameters with *R*^2^ ≥ 0.68 and reduced the correlations between `ksyn_ca` and `intm_bsv` from 0.83 to 0.32. The Peri 5tp protocol further improved recovery (*R*^2^ ≥ 0.83 for all parameters) and reduced the `ksyn_ca`-`intm_bsv` correlation to 0.23. Crucially, the 95% credible interval coverage remained between 94% and 100% across all protocols and parameters, confirming that the posteriors were well-calibrated regardless of sampling intensity and that the shrinkage observed under the single time point protocols is a consequence of a lack of information. The median posterior standard deviation of `intm_bsv` declined progressively from 0.198 to 0.064 log_10_ units (Table 5), matching the progressive removal of the degeneracy.

These results identify three peripheral time points as a clinically efficient compromise: the Peri 3tp protocol recovers 92-96% of the *R*^2^ achieved by the Peri 5tp protocol for four of the five parameters while requiring only three blood draws. Only `intm_bsv`, exhibited a meaningful benefit from five time points (recovering 82% of the Peri 5tp *R*^2^ at three time points). For studies specifically targeting intestinal transit kinetics, the five time point protocol may be warranted, but in general, three time points offer the optimal compromise between identifiability and patient burden.

### Physiological and clinical relevance of five sensitive parameters

Local sensitivity analysis identified five parameters as the primary determinants of fasting peripheral and portal BA concentrations (Fig. 5): the hepatic synthesis rates (`ksyn_ca`, `ksyn_cdca`), the intestinal absorption scaling factor (`lint_abs_bsv`), and the intestinal and colonic transit rates (`intm_bsv`, `colm_bsv`). The identification of these parameters is particularly relevant towards EHC personalization because each could, at least partly, be informed by clinical measurement at the patient- or population-level.

Hepatic BA synthesis can be indirectly assessed through circulating C4, a commonly used marker of BA synthesis, suggesting that synthesis parameters could potentially be individualized using clinical measurements. Intestinal absorption is also clinically relevant because ileal bile acid transporter (IBAT) inhibitors exist on the market and directly reduce BA reabsorption [29]. Data from patients treated with IBAT inhibitors could inform populationor subgroup-specific scaling of the intestinal absorption parameter. Finally, the identification of intestinal transit as an influential process highlights the importance of accounting for variability in gastrointestinal transit and its potential impact on microbial BA metabolism. Although transit time can be measured experimentally via dye or labelled product [30], such measurements may be impractical for routine clinical use. As a first approach, semi-quantitative categories reflecting fast, normal, or slow transit (based on a questionnaire routinely used in clinics) could be used to define subgroup-specific transit parameters.

### Towards personalization of the EHC model for clinical use

Our study was conducted entirely *in silico*: virtual populations were generated from assumed log-normal parameter distributions and observations were synthetic. Therefore, our results demonstrate the performance of the computational pipeline but do not establish parameter identifiability in real patients. Validation using clinical BA data will be an essential and interesting next step. Similarly, the NN surrogate has only been validated within the parameter space covered by the LHS training domain; inference outside these ranges would require additional training and validation with real clinical data.

Beyond personalization for diagnostic purposes, this computational pipeline also establishes the EHC model as a practical tool for mechanistic hypothesis testing. By eliminating the stiff portal and sinusoidal variables and achieving near-instantaneous surrogate evaluations, it becomes computationally feasible to systematically assess which combinations of altered synthesis, feedback regulation, motility, absorption, or microbial transformation reproduce the BA profiles observed in a given disease. Crucially, this approach allows us to determine if and how effectively competing mechanistic hypotheses can be discriminated from a certain set of clinical measurements. The parameter non-identifiabilities identified in our analysis are particularly informative in this regard: they demonstrate that comparable circulating BA profiles can arise from distinct underlying alterations, underscoring that changes in the BA pool cannot be attributed to a single mechanism with certainty based on fasting concentrations alone. Translating this framework to disease states will require extending the surrogate’s training domain across the relevant pathophysiological parameter ranges and validating its predictive accuracy within these domains.

Importantly, our analysis also provides guidance on which additional clinical measurements could improve model personalization. The observed correlations between hepatic synthesis, intestinal motility, and absorption indicate that a single fasting measurement cannot fully distinguish their respective contributions to circulating BA concentrations. For example, similar BA profiles may result from different combinations of synthesis and intestinal transit or absorption. Incorporating measurements that provide complementary information on these processes could therefore improve their individual estimation. Dynamic peripheral BA sampling under fasting and postprandial conditions may capture information on transit and absorption kinetics that is absent from a single fasting measurement. Fecal BA measurements could provide additional information on the fraction of BAs escaping intestinal reab-sorption and thereby help distinguish intestinal motility from absorption, although challenges remain regarding normalization of fecal BA concentrations and their incorporation into ODE models [31].

## Conclusion

Our findings provide a roadmap for personalizing the EHC of BAs model using information from clinically accessible samples. Rather than requiring every physiological parameter to be measured directly, future approaches could combine dynamic multi-time point peripheral BA profiles, fecal measurements, and independent clinical indicators of specific processes with model-based inference. Such integration could progressively constrain patient-specific EHC physiology and ultimately enable the model to translate routinely accessible measurements into individualized estimates of otherwise inaccessible processes and BA profiles. By providing patient-specific, mechanistically grounded BA profiles, this approach will ultimately support the clinical use of BAs as biomarkers for hepatobiliary diseases.

