## Supplementary material for "Temporal blood sampling improves parameter recovery in a model of enterohepatic bile acid circulation using surrogate-accelerated Bayesian inference": S1 File

### Appendix 1: Voronova et al. Enterohepatic Circulation Model ODEs

A description of all parameters and their values can be found in `voronova_parameters.csv`

#### Liver

##### Unconjugated Bile Acids

$$\frac{\mathrm{d}uCA_L}{\mathrm{d}t} = ksyn_{uCA} \cdot (C4 + bln_{C4}) + kupt_{uCA} \cdot hupt_{bsv} \cdot uCA_{SIN} - kref \cdot uCA_L - kconj \cdot fr_{tau}^{conj} \cdot uCA_L - kconj \cdot (1 - fr_{tau}^{conj}) \cdot uCA_L \quad (1)$$

$$\frac{\mathrm{d}uCDCA_L}{\mathrm{d}t} = ksyn_{uCDCA} + kupt_{uCDCA} \cdot uCDCA_{SIN} - kref \cdot hupt_{bsv} \cdot uCDCA_L - kconj \cdot fr_{tau}^{conj} \cdot uCDCA_L - kconj \cdot (1 - fr_{tau}^{conj}) \cdot uCDCA_L \quad (2)$$

$$\frac{\mathrm{d}uDCA_L}{\mathrm{d}t} = kupt_{uDCA} \cdot hupt_{bsv} \cdot uDCA_{SIN} - kref \cdot uDCA_L - kconj \cdot fr_{tau}^{conj} \cdot uDCA_L - kconj \cdot (1 - fr_{tau}^{conj}) \cdot uDCA_L \quad (3)$$

##### Conjugated Bile Acids

$$\frac{\mathrm{d}tCA_L}{\mathrm{d}t} = kconj \cdot fr_{tau}^{conj} \cdot uCA_L + kupt_{tCA} \cdot hupt_{bsv} \cdot tCA_{SIN} - kref \cdot tCA_L - \frac{Q_{ex}}{L} \cdot tCA_L \quad (4)$$

$$\frac{\mathrm{d}gCA_L}{\mathrm{d}t} = kconj \cdot (1 - fr_{tau}^{conj}) \cdot uCA_L + kupt_{gCA} \cdot hupt_{bsv} \cdot gCA_{SIN} - kref \cdot gCA_L - \frac{Q_{ex}}{L} \cdot gCA_L \quad (5)$$

$$\frac{\mathrm{d}tCDCA_L}{\mathrm{d}t} = kconj \cdot fr_{tau}^{conj} \cdot uCDCA_L + kupt_{tCDCA} \cdot hupt_{bsv} \cdot tCDCA_{SIN} - kref \cdot tCDCA_L - \frac{Q_{ex}}{L} \cdot tCDCA_L \quad (6)$$

$$\frac{\mathrm{d}gCDCA_L}{\mathrm{d}t} = kconj \cdot (1 - fr_{tau}^{conj}) \cdot uCDCA_L + kupt_{gCDCA} \cdot hupt_{bsv} \cdot gCDCA_{SIN} - kref \cdot gCDCA_L - \frac{Q_{ex}}{L} \cdot gCDCA_L \quad (7)$$

$$\frac{\mathrm{d}tDCA_L}{\mathrm{d}t} = kconj \cdot fr^{conj}_{tau} \cdot uDCA_L + kupt_{tDCA} \cdot hupt_{bsv} \cdot tDCA_{SIN} - kref \cdot tDCA_L - \frac{Q_{ex}}{L} \cdot tDCA_L \tag{8}$$

$$\frac{\mathrm{d}gDCA_L}{\mathrm{d}t} = kconj \cdot (1 - fr^{conj}_{tau}) \cdot uDCA_L + kupt_{gDCA} \cdot hupt_{bsv} \cdot gDCA_{SIN} - kref \cdot gDCA_L - \frac{Q_{ex}}{L} \cdot gDCA_L \tag{9}$$

### Bile Duct

#### Conjugated Bile Acids

$$\frac{\mathrm{d}tCA_{BD}}{\mathrm{d}t} = \frac{Q_{ex}}{L} \cdot tCA_L - \frac{Q_{sec}}{BD} \cdot fr^{fill}_{GB} \cdot tCA_{BD} - \frac{Q_{sec}}{BD} \cdot (1 - fr^{fill}_{GB}) \cdot tCA_{BD} \tag{10}$$

$$\frac{\mathrm{d}gCA_{BD}}{\mathrm{d}t} = \frac{Q_{ex}}{L} \cdot gCA_L - \frac{Q_{sec}}{BD} \cdot fr^{fill}_{GB} \cdot gCA_{BD} - \frac{Q_{sec}}{BD} \cdot (1 - fr^{fill}_{GB}) \cdot gCA_{BD} \tag{11}$$

$$\frac{\mathrm{d}tCDCA_{BD}}{\mathrm{d}t} = \frac{Q_{ex}}{L} \cdot tCDCA_L - \frac{Q_{sec}}{BD} \cdot fr^{fill}_{GB} \cdot tCDCA_{BD} - \frac{Q_{sec}}{BD} \cdot (1 - fr^{fill}_{GB}) \cdot tCDCA_{BD} \tag{12}$$

$$\frac{\mathrm{d}gCDCA_{BD}}{\mathrm{d}t} = \frac{Q_{ex}}{L} \cdot gCDCA_L - \frac{Q_{sec}}{BD} \cdot fr^{fill}_{GB} \cdot gCDCA_{BD} - \frac{Q_{sec}}{BD} \cdot (1 - fr^{fill}_{GB}) \cdot gCDCA_{BD} \tag{13}$$

$$\frac{\mathrm{d}tDCA_{BD}}{\mathrm{d}t} = \frac{Q_{ex}}{L} \cdot tDCA_L - \frac{Q_{sec}}{BD} \cdot fr^{fill}_{GB} \cdot tDCA_{BD} - \frac{Q_{sec}}{BD} \cdot (1 - fr^{fill}_{GB}) \cdot tDCA_{BD} \tag{14}$$

$$\frac{\mathrm{d}gDCA_{BD}}{\mathrm{d}t} = \frac{Q_{ex}}{L} \cdot gDCA_L - \frac{Q_{sec}}{BD} \cdot fr^{fill}_{GB} \cdot gDCA_{BD} - \frac{Q_{sec}}{BD} \cdot (1 - fr^{fill}_{GB}) \cdot gDCA_{BD} \tag{15}$$

### Gallbladder

#### Conjugated Bile Acids

$$\frac{\mathrm{d}tCA_{GB}}{\mathrm{d}t} = \frac{Q_{sec}}{BD} \cdot fr^{fill}_{GB} \cdot tCA_{BD} - kgbint \cdot gb_{bsv} \cdot tCA_{GB} \cdot a2 \tag{16}$$

$$\frac{\mathrm{d}gCA_{GB}}{\mathrm{d}t} = \frac{Q_{sec}}{BD} \cdot fr^{fill}_{GB} \cdot gCA_{BD} - kgbint \cdot gb_{bsv} \cdot gCA_{GB} \cdot a2 \tag{17}$$

$$\frac{\mathrm{d}tCDCA_{GB}}{\mathrm{d}t} = \frac{Q_{sec}}{BD} \cdot fr^{fill}_{GB} \cdot tCDCA_{BD} - kgbint \cdot gb_{bsv} \cdot tCDCA_{GB} \cdot a2 \tag{18}$$

$$\frac{\mathrm{d}gCDCA_{GB}}{\mathrm{d}t} = \frac{Q_{sec}}{BD} \cdot fr^{fill}_{GB} \cdot gCDCA_{BD} - kgbint \cdot gb_{bsv} \cdot gCDCA_{GB} \cdot a2 \tag{19}$$

$$\frac{\mathrm{d}tDCA_{GB}}{\mathrm{d}t} = \frac{Q_{sec}}{BD} \cdot fr_{GB}^{fill} \cdot tDCA_{BD} - kgbint \cdot gb_{bsv} \cdot tDCA_{GB} \cdot a2 \tag{20}$$

$$\frac{\mathrm{d}gDCA_{GB}}{\mathrm{d}t} = \frac{Q_{sec}}{BD} \cdot fr_{GB}^{fill} \cdot gDCA_{BD} - kgbint \cdot gb_{bsv} \cdot gDCA_{GB} \cdot a2 \tag{21}$$

### Upper Intestine

#### Conjugated Bile Acids

$$\begin{aligned}\frac{\mathrm{d}tCA_{UINT}}{\mathrm{d}t} = kgbint \cdot gb_{bsv} \cdot tCA_{GB} \cdot a2 + \frac{Q_{sec}}{BD} \cdot fr_{GB}^{fill} \cdot tCA_{BD} \\ - ktr_{INTINT} \cdot (1 + a1 \cdot gm) \cdot intm_{bsv} \cdot tCA_{UINT}\end{aligned} \tag{22}$$

$$\begin{aligned}\frac{\mathrm{d}gCA_{UINT}}{\mathrm{d}t} = kgbint \cdot gb_{bsv} \cdot gCA_{GB} \cdot a2 + \frac{Q_{sec}}{BD} \cdot fr_{GB}^{fill} \cdot gCA_{BD} \\ - ktr_{INTINT} \cdot (1 + a1 \cdot gm) \cdot intm_{bsv} \cdot gCA_{UINT}\end{aligned} \tag{23}$$

$$\begin{aligned}\frac{\mathrm{d}tCDCA_{UINT}}{\mathrm{d}t} = kgbint \cdot gb_{bsv} \cdot tCDCA_{GB} \cdot a2 + \frac{Q_{sec}}{BD} \cdot fr_{GB}^{fill} \cdot tCDCA_{BD} \\ - ktr_{INTINT} \cdot (1 + a1 \cdot gm) \cdot intm_{bsv} \cdot tCDCA_{UINT}\end{aligned} \tag{24}$$

$$\begin{aligned}\frac{\mathrm{d}gCDCA_{UINT}}{\mathrm{d}t} = kgbint \cdot gb_{bsv} \cdot gCDCA_{GB} \cdot a2 + \frac{Q_{sec}}{BD} \cdot fr_{GB}^{fill} \cdot gCDCA_{BD} \\ - kintabs_{gCDCA} \cdot uintabs_{bsv} \cdot gCDCA_{UINT} \\ - ktr_{INTINT} \cdot (1 + a1 \cdot gm) \cdot intm_{bsv} \cdot gCDCA_{UINT}\end{aligned} \tag{25}$$

$$\begin{aligned}\frac{\mathrm{d}tDCA_{UINT}}{\mathrm{d}t} = kgbint \cdot gb_{bsv} \cdot tDCA_{GB} \cdot a2 + \frac{Q_{sec}}{BD} \cdot fr_{GB}^{fill} \cdot tDCA_{BD} \\ - ktr_{INTINT} \cdot (1 + a1 \cdot gm) \cdot intm_{bsv} \cdot tDCA_{UINT}\end{aligned} \tag{26}$$

$$\begin{aligned}\frac{\mathrm{d}gDCA_{UINT}}{\mathrm{d}t} = kgbint \cdot gb_{bsv} \cdot gDCA_{GB} \cdot a2 + \frac{Q_{sec}}{BD} \cdot fr_{GB}^{fill} \cdot gDCA_{BD} \\ - kintabs_{gDCA} \cdot uintabs_{bsv} \cdot gDCA_{UINT} \\ - ktr_{INTINT} \cdot (1 + a1 \cdot gm) \cdot intm_{bsv} \cdot gDCA_{UINT}\end{aligned} \tag{27}$$

### Lower Intestine

#### Conjugated Bile Acids

$$\begin{aligned}\frac{\mathrm{d}tCA_{LINT}}{\mathrm{d}t} = ktr_{INTINT} \cdot (1+a1 \cdot gm) \cdot intm_{bsv} \cdot tCA_{UINT} - ktr_{INTCOL} \cdot (1+a1 \cdot gm) \cdot intm_{bsv} \cdot tCA_{LINT} \\ - kdcj_{tba_{LINT}} \cdot lint_{dcj} \cdot (1 - a1 \cdot tmbt) \cdot lintdcj_{bsv} \cdot tCA_{LINT} \\ - klintabs_{tCA} \cdot lintabs_{bsv} \cdot tCA_{LINT}\end{aligned} \tag{28}$$

$$\frac{\mathrm{d}gCA_{LINT}}{\mathrm{d}t} = ktr_{INTINT}\cdot(1+a1\cdot gm)\cdot intm_{bsv}\cdot gCA_{UINT} - ktr_{INTCOL}\cdot(1+a1\cdot gm)\cdot intm_{bsv}\cdot gCA_{LINT} - kdcj_{gba_{LINT}}\cdot lint_{dcj}\cdot(1-a1\cdot tmbt)\cdot lintdcj_{bsv}\cdot gCA_{LINT} - klintabs_{gCA}\cdot lintabs_{bsv}\cdot gCA_{LINT} \tag{29}$$

$$\frac{\mathrm{d}tCDCA_{LINT}}{\mathrm{d}t} = ktr_{INTINT}\cdot(1+a1\cdot gm)\cdot intm_{bsv}\cdot tCDCA_{UINT} - ktr_{INTCOL}\cdot(1+a1\cdot gm)\cdot intm_{bsv}\cdot tCDCA_{LINT} - kdcj_{tba_{LINT}}\cdot lint_{dcj}\cdot(1-a1\cdot tmbt)\cdot lintdcj_{bsv}\cdot tCDCA_{LINT} - klintabs_{tCDCA}\cdot lintabs_{bsv}\cdot tCDCA_{LINT} \tag{30}$$

$$\frac{\mathrm{d}gCDCA_{LINT}}{\mathrm{d}t} = ktr_{INTINT}\cdot(1+a1\cdot gm)\cdot intm_{bsv}\cdot gCDCA_{UINT} - ktr_{INTCOL}\cdot(1+a1\cdot gm)\cdot intm_{bsv}\cdot gCDCA_{LINT} - kdcj_{gba_{LINT}}\cdot lint_{dcj}\cdot(1-a1\cdot tmbt)\cdot lintdcj_{bsv}\cdot gCDCA_{LINT} - klintabs_{gCDCA}\cdot lintabs_{bsv}\cdot gCDCA_{LINT} \tag{31}$$

$$\frac{\mathrm{d}tDCA_{LINT}}{\mathrm{d}t} = ktr_{INTINT}\cdot(1+a1\cdot gm)\cdot intm_{bsv}\cdot tDCA_{UINT} - ktr_{INTCOL}\cdot(1+a1\cdot gm)\cdot intm_{bsv}\cdot tDCA_{LINT} - kdcj_{tba_{LINT}}\cdot lint_{dcj}\cdot(1-a1\cdot tmbt)\cdot lintdcj_{bsv}\cdot tDCA_{LINT} - klintabs_{tDCA}\cdot lintabs_{bsv}\cdot tDCA_{LINT} \tag{32}$$

$$\frac{\mathrm{d}gDCA_{LINT}}{\mathrm{d}t} = ktr_{INTINT}\cdot(1+a1\cdot gm)\cdot intm_{bsv}\cdot gDCA_{UINT} - ktr_{INTCOL}\cdot(1+a1\cdot gm)\cdot intm_{bsv}\cdot gDCA_{LINT} - kdcj_{gba_{LINT}}\cdot lint_{dcj}\cdot(1-a1\cdot tmbt)\cdot lintdcj_{bsv}\cdot gDCA_{LINT} - klintabs_{gDCA}\cdot lintabs_{bsv}\cdot gDCA_{LINT} \tag{33}$$

#### Unconjugated Bile Acids

$$\frac{\mathrm{d}uCA_{LINT}}{\mathrm{d}t} = kdcj_{tba_{LINT}}\cdot lint_{dcj}\cdot(1-a1\cdot tmbt)\cdot lintdcj_{bsv}\cdot tCA_{LINT} + kdcj_{gba_{LINT}}\cdot lint_{dcj}\cdot(1-a1\cdot tmbt)\cdot lintdcj_{bsv}\cdot gCA_{LINT} - ktr_{INTCOL}\cdot(1+a1\cdot gm)\cdot intm_{bsv}\cdot uCA_{LINT} - klintabs_{uCA}\cdot lintabs_{bsv}\cdot uCA_{LINT} \tag{34}$$

$$\frac{\mathrm{d}uCDCA_{LINT}}{\mathrm{d}t} = kdcj_{tba_{LINT}}\cdot lint_{dcj}\cdot(1-a1\cdot tmbt)\cdot lintdcj_{bsv}\cdot tCDCA_{LINT} + kdcj_{gba_{LINT}}\cdot lint_{dcj}\cdot(1-a1\cdot tmbt)\cdot lintdcj_{bsv}\cdot gCDCA_{LINT} - ktr_{INTCOL}\cdot(1+a1\cdot gm)\cdot intm_{bsv}\cdot uCDCA_{LINT} - klintabs_{uCDCA}\cdot lintabs_{bsv}\cdot uCDCA_{LINT} \tag{35}$$

$$\frac{\mathrm{d}uDCA_{LINT}}{\mathrm{d}t} = kdcj_{tba_{LINT}}\cdot lint_{dcj}\cdot(1-a1\cdot tmbt)\cdot lintdcj_{bsv}\cdot tDCA_{LINT} + kdcj_{gba_{LINT}}\cdot lint_{dcj}\cdot(1-a1\cdot tmbt)\cdot lintdcj_{bsv}\cdot gDCA_{LINT} - ktr_{INTCOL}\cdot(1+a1\cdot gm)\cdot intm_{bsv}\cdot uDCA_{LINT} - klintabs_{uDCA}\cdot lintabs_{bsv}\cdot uDCA_{LINT} \tag{36}$$

### Colon

#### Conjugated Bile Acids

$$\frac{\mathrm{d}tCA_{COL}}{\mathrm{d}t} = ktr_{INTCOL} \cdot (1 + a1 \cdot gm) \cdot intm_{bsv} \cdot tCA_{LINT} - kcoldcj \cdot coldcj_{bsv} \cdot tCA_{COL} \quad (37)$$

$$\frac{\mathrm{d}gCA_{COL}}{\mathrm{d}t} = ktr_{INTCOL} \cdot (1 + a1 \cdot gm) \cdot intm_{bsv} \cdot gCA_{LINT} - kcoldcj \cdot coldcj_{bsv} \cdot gCA_{COL} \quad (38)$$

$$\frac{\mathrm{d}tCDCA_{COL}}{\mathrm{d}t} = ktr_{INTCOL} \cdot (1 + a1 \cdot gm) \cdot intm_{bsv} \cdot tCDCA_{LINT} - kcoldcj \cdot coldcj_{bsv} \cdot tCDCA_{COL} \quad (39)$$

$$\frac{\mathrm{d}gCDCA_{COL}}{\mathrm{d}t} = ktr_{INTCOL} \cdot (1 + a1 \cdot gm) \cdot intm_{bsv} \cdot gCDCA_{LINT} - kcoldcj \cdot coldcj_{bsv} \cdot gCDCA_{COL} \quad (40)$$

$$\frac{\mathrm{d}tDCA_{COL}}{\mathrm{d}t} = ktr_{INTCOL} \cdot (1 + a1 \cdot gm) \cdot intm_{bsv} \cdot tDCA_{LINT} - kcoldcj \cdot coldcj_{bsv} \cdot tDCA_{COL} \quad (41)$$

$$\frac{\mathrm{d}gDCA_{COL}}{\mathrm{d}t} = ktr_{INTCOL} \cdot (1 + a1 \cdot gm) \cdot intm_{bsv} \cdot gDCA_{LINT} - kcoldcj \cdot coldcj_{bsv} \cdot gDCA_{COL} \quad (42)$$

#### Unconjugated Bile Acids

$$\frac{\mathrm{d}uCA_{COL}}{\mathrm{d}t} = ktr_{INTCOL} \cdot (1 + a1 \cdot gm) \cdot intm_{bsv} \cdot uCA_{LINT} + kcoldcj \cdot coldcj_{bsv} \cdot tCA_{COL} + kcoldcj \cdot coldcj_{bsv} \cdot gCA_{COL} - ksyn_{DCA} \cdot mbt_{bsv} \cdot uCA_{COL} - kcolel \cdot colm_{bsv} \cdot uCA_{COL} - kcolabs_{CA} \cdot colabs_{bsv} \cdot uCA_{COL} \quad (43)$$

$$\frac{\mathrm{d}uCDCA_{COL}}{\mathrm{d}t} = ktr_{INTCOL} \cdot (1 + a1 \cdot gm) \cdot intm_{bsv} \cdot uCDCA_{LINT} + kcoldcj \cdot coldcj_{bsv} \cdot tCDCA_{COL} + kcoldcj \cdot coldcj_{bsv} \cdot gCDCA_{COL} - kel_{CDCA} \cdot mbt_{bsv} \cdot uCDCA_{COL} - kcolel \cdot colm_{bsv} \cdot uCDCA_{COL} - kcolabs_{CDCA} \cdot colabs_{bsv} \cdot uCDCA_{COL} \quad (44)$$

$$\frac{\mathrm{d}uDCAin_{COL}}{\mathrm{d}t} = ksyn_{DCA} \cdot mbt_{bsv} \cdot uCA_{COL} - ksol_{DCA} \cdot uDCAin_{COL} - kcolel \cdot colm_{bsv} \cdot uDCAin_{COL} \quad (45)$$

$$\frac{\mathrm{d}uDCA_{COL}}{\mathrm{d}t} = ktr_{INTCOL} \cdot (1 + a1 \cdot gm) \cdot intm_{bsv} \cdot uDCA_{LINT} + kcoldcj \cdot coldcj_{bsv} \cdot tDCA_{COL} + kcoldcj \cdot coldcj_{bsv} \cdot gDCA_{COL} + ksol_{DCA} \cdot uDCAin_{COL} - kcolel \cdot colm_{bsv} \cdot uDCA_{COL} - kcolabs_{DCA} \cdot colabs_{bsv} \cdot uDCA_{COL} \quad (46)$$

### Portal Vein

#### Unconjugated Bile Acids

$$\frac{\mathrm{d}uCA_{PV}}{\mathrm{d}t} = \frac{Q_{MA}}{SYS} \cdot (1 + a1 \cdot pc) \cdot uCA_{SYS} + klintabs_{uCA} \cdot lintabs_{bsv} \cdot uCA_{LINT} + kcolabs_{CA} \cdot colabs_{bsv} \cdot uCA_{COL} - \frac{Q_{PV}}{PV} \cdot (1 + a1 \cdot pc) \cdot uCA_{PV} \quad (47)$$

$$\frac{\mathrm{d}uCDCA_{PV}}{\mathrm{d}t} = \frac{Q_{MA}}{SYS} \cdot (1 + a1 \cdot pc) \cdot uCDCA_{SYS} + klintabs_{uCDCADCA} \cdot lintabs_{bsv} \cdot uCDCA_{LINT} + kcolabs_{CDCA} \cdot colabs_{bsv} \cdot uCDCA_{COL} - \frac{Q_{PV}}{PV} \cdot (1 + a1 \cdot pc) \cdot uCDCA_{PV} \quad (48)$$

$$\frac{\mathrm{d}uDCA_{PV}}{\mathrm{d}t} = \frac{Q_{MA}}{SYS} \cdot (1 + a1 \cdot pc) \cdot uDCA_{SYS} + klintabs_{uCDCADCA} \cdot lintabs_{bsv} \cdot uDCA_{LINT} + kcolabs_{DCA} \cdot colabs_{bsv} \cdot uDCA_{COL} - \frac{Q_{PV}}{PV} \cdot (1 + a1 \cdot pc) \cdot uDCA_{PV} \quad (49)$$

#### Conjugated Bile Acids

$$\frac{\mathrm{d}tCA_{PV}}{\mathrm{d}t} = \frac{Q_{MA}}{SYS} \cdot (1 + a1 \cdot pc) \cdot tCA_{SYS} + klintabs_{tCA} \cdot lintabs_{bsv} \cdot tCA_{LINT} - \frac{Q_{PV}}{PV} \cdot (1 + a1 \cdot pc) \cdot tCA_{PV} \quad (50)$$

$$\frac{\mathrm{d}gCA_{PV}}{\mathrm{d}t} = \frac{Q_{MA}}{SYS} \cdot (1 + a1 \cdot pc) \cdot gCA_{SYS} + klintabs_{gCA} \cdot lintabs_{bsv} \cdot gCA_{LINT} - \frac{Q_{PV}}{PV} \cdot (1 + a1 \cdot pc) \cdot gCA_{PV} \quad (51)$$

$$\frac{\mathrm{d}tCDCA_{PV}}{\mathrm{d}t} = \frac{Q_{MA}}{SYS} \cdot (1 + a1 \cdot pc) \cdot tCDCA_{SYS} + klintabs_{tCDCA} \cdot lintabs_{bsv} \cdot tCDCA_{LINT} - \frac{Q_{PV}}{PV} \cdot (1 + a1 \cdot pc) \cdot tCDCA_{PV} \quad (52)$$

$$\frac{\mathrm{d}gCDCA_{PV}}{\mathrm{d}t} = \frac{Q_{MA}}{SYS} \cdot (1 + a1 \cdot pc) \cdot gCDCA_{SYS} + kintabs_{tCDCA} \cdot uintabs_{bsv} \cdot gCDCA_{UINT} + klintabs_{gCDCA} \cdot lintabs_{bsv} \cdot gCDCA_{LINT} - \frac{Q_{PV}}{PV} \cdot (1 + a1 \cdot pc) \cdot gCDCA_{PV} \quad (53)$$

$$\frac{\mathrm{d}tDCA_{PV}}{\mathrm{d}t} = \frac{Q_{MA}}{SYS} \cdot (1 + a1 \cdot pc) \cdot tDCA_{SYS} + klintabs_{tDCA} \cdot lintabs_{bsv} \cdot tDCA_{LINT} - \frac{Q_{PV}}{PV} \cdot (1 + a1 \cdot pc) \cdot tDCA_{PV} \quad (54)$$

$$\frac{\mathrm{d}gDCA_{PV}}{\mathrm{d}t} = \frac{Q_{MA}}{SYS} \cdot (1 + a1 \cdot pc) \cdot gDCA_{SYS} + kintabs_{gDCA} \cdot uintabs_{bsv} \cdot gDCA_{UINT} + klintabs_{gDCA} \cdot lintabs_{bsv} \cdot gDCA_{LINT} - \frac{Q_{PV}}{PV} \cdot (1 + a1 \cdot pc) \cdot gDCA_{PV} \quad (55)$$

#### Sinusoidal Space

##### Unconjugated Bile Acids

$$\frac{\mathrm{d}uCA_{SIN}}{\mathrm{d}t} = \frac{Q_{PV}}{PV} \cdot (1 + a1 \cdot pc) \cdot uCA_{PV} + \frac{Q_{HA}}{SYS} \cdot (1 + a1 \cdot pc) \cdot uCA_{SYS} + kref \cdot uCA_L - \frac{Q_{HV}}{SIN} \cdot (1 + a1 \cdot pc) \cdot uCA_{SIN} - kupt_{uCA} \cdot hupt_{bsv} \cdot uCA_{SIN} \quad (56)$$

$$\frac{\mathrm{d}uCDCA_{SIN}}{\mathrm{d}t} = \frac{Q_{PV}}{PV} \cdot (1 + a1 \cdot pc) \cdot uCDCA_{PV} + \frac{Q_{HA}}{SYS} \cdot (1 + a1 \cdot pc) \cdot uCDCA_{SYS} + kref \cdot uCDCA_L - \frac{Q_{HV}}{SIN} \cdot (1 + a1 \cdot pc) \cdot uCDCA_{SIN} - kupt_{uCDCA} \cdot hupt_{bsv} \cdot uCDCA_{SIN} \quad (57)$$

$$\frac{\mathrm{d}uDCA_{SIN}}{\mathrm{d}t} = \frac{Q_{PV}}{PV} \cdot (1 + a1 \cdot pc) \cdot uDCA_{PV} + \frac{Q_{HA}}{SYS} \cdot (1 + a1 \cdot pc) \cdot uDCA_{SYS} + kref \cdot uDCA_L - \frac{Q_{HV}}{SIN} \cdot (1 + a1 \cdot pc) \cdot uDCA_{SIN} - kupt_{uDCA} \cdot hupt_{bsv} \cdot uDCA_{SIN} \quad (58)$$

##### Conjugated Bile Acids

$$\frac{\mathrm{d}tCA_{SIN}}{\mathrm{d}t} = \frac{Q_{PV}}{PV} \cdot (1 + a1 \cdot pc) \cdot tCA_{PV} + \frac{Q_{HA}}{SYS} \cdot (1 + a1 \cdot pc) \cdot tCA_{SYS} + kref \cdot tCA_L - \frac{Q_{HV}}{SIN} \cdot (1 + a1 \cdot pc) \cdot tCA_{SIN} - kupt_{tCA} \cdot hupt_{bsv} \cdot tCA_{SIN} \quad (59)$$

$$\frac{\mathrm{d}gCA_{SIN}}{\mathrm{d}t} = \frac{Q_{PV}}{PV} \cdot (1 + a1 \cdot pc) \cdot gCA_{PV} + \frac{Q_{HA}}{SYS} \cdot (1 + a1 \cdot pc) \cdot gCA_{SYS} + kref \cdot gCA_L - \frac{Q_{HV}}{SIN} \cdot (1 + a1 \cdot pc) \cdot gCA_{SIN} - kupt_{gCA} \cdot hupt_{bsv} \cdot gCA_{SIN} \quad (60)$$

$$\frac{\mathrm{d}tCDCA_{SIN}}{\mathrm{d}t} = \frac{Q_{PV}}{PV} \cdot (1 + a1 \cdot pc) \cdot tCDCA_{PV} + \frac{Q_{HA}}{SYS} \cdot (1 + a1 \cdot pc) \cdot tCDCA_{SYS} + kref \cdot tCDCA_L - \frac{Q_{HV}}{SIN} \cdot (1 + a1 \cdot pc) \cdot tCDCA_{SIN} - kupt_{tCDCA} \cdot hupt_{bsv} \cdot tCDCA_{SIN} \quad (61)$$

$$\frac{\mathrm{d}gCDCA_{SIN}}{\mathrm{d}t} = \frac{Q_{PV}}{PV} \cdot (1 + a1 \cdot pc) \cdot gCDCA_{PV} + \frac{Q_{HA}}{SYS} \cdot (1 + a1 \cdot pc) \cdot gCDCA_{SYS} + kref \cdot gCDCA_L - \frac{Q_{HV}}{SIN} \cdot (1 + a1 \cdot pc) \cdot gCDCA_{SIN} - kupt_{gCDCA} \cdot hupt_{bsv} \cdot gCDCA_{SIN} \quad (62)$$

$$\frac{\mathrm{d}tDCA_{SIN}}{\mathrm{d}t} = \frac{Q_{PV}}{PV} \cdot (1 + a1 \cdot pc) \cdot tDCA_{PV} + \frac{Q_{HA}}{SYS} \cdot (1 + a1 \cdot pc) \cdot tDCA_{SYS} + kref \cdot tDCA_L - \frac{Q_{HV}}{SIN} \cdot (1 + a1 \cdot pc) \cdot tDCA_{SIN} - kupt_{tDCA} \cdot hupt_{bsv} \cdot tDCA_{SIN} \quad (63)$$

$$\frac{\mathrm{d}gDCA_{SIN}}{\mathrm{d}t} = \frac{Q_{PV}}{PV} \cdot (1 + a1 \cdot pc) \cdot gDCA_{PV} + \frac{Q_{HA}}{SYS} \cdot (1 + a1 \cdot pc) \cdot gDCA_{SYS} + kref \cdot gDCA_L - \frac{Q_{HV}}{SIN} \cdot (1 + a1 \cdot pc) \cdot gDCA_{SIN} - kupt_{gDCA} \cdot hupt_{bsv} \cdot gDCA_{SIN} \quad (64)$$

#### Systemic Circulation

##### Unconjugated Bile Acids

$$\frac{\mathrm{d}uCA_{SYS}}{\mathrm{d}t} = \frac{Q_{HV}}{SIN} \cdot (1 + a1 \cdot pc) \cdot uCA_{SIN} - \frac{Q_{HA}}{SYS} \cdot (1 + a1 \cdot pc) \cdot uCA_{SYS} - \frac{Q_{MA}}{SYS} \cdot (1 + a1 \cdot pc) \cdot uCA_{SYS} \quad (65)$$

$$\frac{\mathrm{d}uCDCA_{SYS}}{\mathrm{d}t} = \frac{Q_{HV}}{SIN} \cdot (1 + a1 \cdot pc) \cdot uCDCA_{SIN} - \frac{Q_{HA}}{SYS} \cdot (1 + a1 \cdot pc) \cdot uCDCA_{SYS} - \frac{Q_{MA}}{SYS} \cdot (1 + a1 \cdot pc) \cdot uCDCA_{SYS} \quad (66)$$

$$\frac{\mathrm{d}uDCA_{SYS}}{\mathrm{d}t} = \frac{Q_{HV}}{SIN} \cdot (1 + a1 \cdot pc) \cdot uDCA_{SIN} - \frac{Q_{HA}}{SYS} \cdot (1 + a1 \cdot pc) \cdot uDCA_{SYS} - \frac{Q_{MA}}{SYS} \cdot (1 + a1 \cdot pc) \cdot uDCA_{SYS} \quad (67)$$

##### Conjugated Bile Acids

$$\frac{\mathrm{d}tCA_{SYS}}{\mathrm{d}t} = \frac{Q_{HV}}{SIN} \cdot (1 + a1 \cdot pc) \cdot tCA_{SIN} - \frac{Q_{HA}}{SYS} \cdot (1 + a1 \cdot pc) \cdot tCA_{SYS} - \frac{Q_{MA}}{SYS} \cdot (1 + a1 \cdot pc) \cdot tCA_{SYS} \quad (68)$$

$$\frac{\mathrm{d}gCA_{SYS}}{\mathrm{d}t} = \frac{Q_{HV}}{SIN} \cdot (1 + a1 \cdot pc) \cdot gCA_{SIN} - \frac{Q_{HA}}{SYS} \cdot (1 + a1 \cdot pc) \cdot gCA_{SYS} - \frac{Q_{MA}}{SYS} \cdot (1 + a1 \cdot pc) \cdot gCA_{SYS} \quad (69)$$

$$\frac{\mathrm{d}tCDCA_{SYS}}{\mathrm{d}t} = \frac{Q_{HV}}{SIN} \cdot (1 + a1 \cdot pc) \cdot tCDCA_{SIN} - \frac{Q_{HA}}{SYS} \cdot (1 + a1 \cdot pc) \cdot tCDCA_{SYS} - \frac{Q_{MA}}{SYS} \cdot (1 + a1 \cdot pc) \cdot tCDCA_{SYS} \quad (70)$$

$$\frac{\mathrm{d}gCDCA_{SYS}}{\mathrm{d}t} = \frac{Q_{HV}}{SIN} \cdot (1 + a1 \cdot pc) \cdot gCDCA_{SIN} - \frac{Q_{HA}}{SYS} \cdot (1 + a1 \cdot pc) \cdot gCDCA_{SYS} - \frac{Q_{MA}}{SYS} \cdot (1 + a1 \cdot pc) \cdot gCDCA_{SYS} \quad (71)$$

$$\frac{\mathrm{d}tDCA_{SYS}}{\mathrm{d}t} = \frac{Q_{HV}}{SIN} \cdot (1 + a1 \cdot pc) \cdot tDCA_{SIN} - \frac{Q_{HA}}{SYS} \cdot (1 + a1 \cdot pc) \cdot tDCA_{SYS} - \frac{Q_{MA}}{SYS} \cdot (1 + a1 \cdot pc) \cdot tDCA_{SYS} \quad (72)$$

$$\frac{\mathrm{d}gDCA_{SYS}}{\mathrm{d}t} = \frac{Q_{HV}}{SIN} \cdot (1 + a1 \cdot pc) \cdot gDCA_{SIN} - \frac{Q_{HA}}{SYS} \cdot (1 + a1 \cdot pc) \cdot gDCA_{SYS} - \frac{Q_{MA}}{SYS} \cdot (1 + a1 \cdot pc) \cdot gDCA_{SYS} \quad (73)$$

#### Bile Acid Synthesis Feedback Loop

$$\frac{\mathrm{d}FGF19}{\mathrm{d}t} = kdel \cdot (FXRa - FGF19) \tag{74}$$

where

$$FXR_a = Fmax_{FXR} \cdot \frac{\frac{1}{EC50_{FXR}} \sum_i \frac{[BA]_{i,LINT}}{fFXR_i}}{\frac{1}{EC50_{FXR}} \sum_i \frac{[BA]_{i,LINT}}{fFXR_i} + 1}$$

$$FXR_a = Fmax_{FXR} \cdot \frac{\frac{\frac{tCDCA_{LINT}+gCDCA_{LINT}+tDCA_{LINT}+gDCA_{LINT}}{fFXR_{cCDCADCA}} + \frac{tCA_{LINT}+gCA_{LINT}}{fFXR_{cCA}} + \frac{uCDCA_{LINT}}{fFXR_{uCDCA}} + \frac{uDCA_{LINT}}{fFXR_{uDCA}}}{EC50_{FXR}}}{\frac{\frac{tCDCA_{LINT}+gCDCA_{LINT}+tDCA_{LINT}+gDCA_{LINT}}{fFXR_{cCDCADCA}} + \frac{tCA_{LINT}+gCA_{LINT}}{fFXR_{cCA}} + \frac{uCDCA_{LINT}}{fFXR_{uCDCA}} + \frac{uDCA_{LINT}}{fFXR_{uDCA}}}{EC50_{FXR}} + 1}$$

In Eq. 1,

$$C4 = \exp((1 - FGF19) \cdot a_{ba_{FGF19}})$$
