## Supplementary material for "Temporal blood sampling improves parameter recovery in a model of enterohepatic bile acid circulation using surrogate-accelerated Bayesian inference": S3 File

### Appendix 3: Inhibitory Hill Function Derivation

The activation of the nuclear receptor $FXR$ by bile acids produces $FGF19$. This binds to and activates $FGFR4$, which leads to the suppression of CYP7A1 in the liver, and thus less $7\alpha$-hydroxy-4-cholesten-3-one ($C4$) production, a precursor of bile acids.

#### Original Model

$$\frac{\mathrm{d}FGF19}{\mathrm{d}t} = kdel \cdot (FXRa - FGF19) \tag{1}$$

where

$$FXR_a = Fmax_{FXR} \cdot \frac{\frac{1}{EC50_{FXR}} \sum_i \frac{[BA]_{i,LINT}}{fFXR_i}}{\frac{1}{EC50_{FXR}} \sum_i \frac{[BA]_{i,LINT}}{fFXR_i} + 1}$$

$$FXR_a = Fmax_{FXR} \cdot \frac{\frac{\frac{tCDCA_{LINT}+gCDCA_{LINT}+tDCA_{LINT}+gDCA_{LINT}}{fFXR_{cCDCADCA}} + \frac{tCA_{LINT}+gCA_{LINT}}{fFXR_{cCA}} + \frac{uCDCA_{LINT}}{fFXR_{uCDCA}} + \frac{uDCA_{LINT}}{fFXR_{uDCA}}}{EC50_{FXR}}}{\frac{\frac{tCDCA_{LINT}+gCDCA_{LINT}+tDCA_{LINT}+gDCA_{LINT}}{fFXR_{cCDCADCA}} + \frac{tCA_{LINT}+gCA_{LINT}}{fFXR_{cCA}} + \frac{uCDCA_{LINT}}{fFXR_{uCDCA}} + \frac{uDCA_{LINT}}{fFXR_{uDCA}}}{EC50_{FXR}} + 1}$$

The amount of $FGF19$ is substituted in the algebraic equation for the amount of C4 at any given time:

$$C4 = \exp((1 - FGF19) \cdot a_{ba_{FGF19}}) \tag{2}$$

which in turn is substituted into the ODE for $uCA$ in the liver ($uCA_L$):

$$\begin{aligned} \frac{\mathrm{d}uCA_L}{\mathrm{d}t} &= ksyn_{uCA} \cdot (C4 + bln_{C4}) + kupt_{uCA} \cdot hupt_{bsv} \cdot uCA_{SIN} \\ &\quad - kref \cdot uCA_L - kconj \cdot fr_{tau}^{conj} \cdot uCA_L - kconj \cdot (1 - fr_{tau}^{conj}) \cdot uCA_L \end{aligned} \tag{3}$$

where $bln_{C4}$ is a constant baseline offset calculated to ensure that the area under the curve (AUC) of $C4$ over a 24-hour period at steady state is equal to 1.

#### Modifications

A traditional inhibitory Hill function will be used instead:

$$E = E_{max} \cdot \frac{IC_{50}^n}{IC_{50}^n + [I]^n} \tag{4}$$

where $E$ represents the observed effect, $E_{max}$ is the maximum possible effect in the absence of inhibition, $[I]$ is the concentration of the inhibitor, $IC_{50}$ is the concentration of the inhibitor that reduces the effect by half, and $n$ is the Hill coefficient dictating the cooperativity of the binding.

In the context of this model the biological effect ($E$) corresponds to the C4 synthesis multiplier, and the inhibitor ($[I]$) is the normalized FGF19 signal. Assuming simple 1-to-1, non-cooperative binding between the ligand and the regulatory target, the Hill coefficient is defined as $n = 1$. This simplifies the regulatory feedback loop to a standard $I_{max}$ function:

$$\boldsymbol{C4 = C4_{max} \cdot \frac{IC_{50}}{IC_{50} + FGF19}} \tag{5}$$

The kinetic parameters were algebraically derived to ensure the modified model conserves some behaviour of the original system, specifically the same absolute maximum synthesis and steady-state baseline.

The original exponential feedback equation is defined as:

$$C4_{exp} = \text{exp}((1 - FGF19) \cdot a_{ba_{FGF19}}) \tag{6}$$

Theoretically, maximum synthesis occurs in the absence of negative feedback($FGF19 = 0$). Evaluating the original exponential model at this boundary provides the maximum value for synthesis. Setting the modified equation equal to this value yields $C4_{max}$:

$$C4_{max} = \text{exp}(a_{ba_{FGF19}}) \tag{7}$$

Furthermore, physiological baseline steady-state occurs when the normalized $FGF19$ signal is exactly $1$. Evaluating the original model at this baseline yields a synthesis multiplier of $1$ (since exp$(0) = 1$). To maintain the baseline mass balance, the modified equation must also equal $1$ when $FGF19 = 1$:

$$1 = C4_{max} \cdot \frac{IC_{50}}{IC_{50} + 1} \tag{8}$$

Substituting $C4_{max}$ into this equation allows for the derivation of $IC_{50}$:

$$\begin{aligned} 1 &= \text{exp}(a_{ba_{FGF19}}) \cdot \frac{IC_{50}}{IC_{50} + 1} \\ IC_{50} + 1 &= \text{exp}(a_{ba_{FGF19}}) \cdot IC_{50} \\ 1 &= IC_{50} \cdot (\text{exp}(a_{ba_{FGF19}}) - 1) \\ \boldsymbol{IC_{50}} &= \boldsymbol{\frac{1}{\text{exp}(a_{ba_{FGF19}}) - 1}} \end{aligned}$$

By substituting these algebraically derived parameters back into the modified model, the new function maps the original arbitrary decay constant ($a_{ba_{FGF19}}$) into standard pharmacological metrics and preserves the original steady-state boundaries.

Because the new $I_{max}$ function intrinsically captures baseline synthesis when $FGF19 = 1$, the fixed synthesis parameter ($bln_{c4}$) was removed from the ODE for $uCA_L$:

$$\begin{aligned} \frac{\text{d}uCA_L}{\text{d}t} = &\ ksyn_{uCA} \cdot (\boldsymbol{C4}) + kupt_{uCA} \cdot hupt_{bsv} \cdot uCA_{SIN} \\ &- kref \cdot uCA_L - kconj \cdot fr_{tau}^{conj} \cdot uCA_L - kconj \cdot (1 - fr_{tau}^{conj}) \cdot uCA_L \end{aligned} \tag{9}$$

By eliminating this fixed offset, the modified model relies solely on the dynamic, receptor-mediated $I_{max}$ function to drive and regulate basal bile acid synthesis.
