## Supplementary material for "Temporal blood sampling improves parameter recovery in a model of enterohepatic bile acid circulation using surrogate-accelerated Bayesian inference": S4 File

### Appendix 4: Quasi-Steady State Approximation Equations

A description of all parameters and their values can be found in `voronova_parameters.csv`

#### Portal Vein

##### Unconjugated Bile Acids

$$\frac{\mathrm{d}uCA_{PV}}{\mathrm{d}t} = \frac{Q_{MA}}{SYS} \cdot (1 + a1 \cdot pc) \cdot uCA_{SYS} + klintabs_{uCA} \cdot lintabs_{bsv} \cdot uCA_{LINT} + kcolabs_{CA} \cdot colabs_{bsv} \cdot uCA_{COL} - \frac{Q_{PV}}{PV} \cdot (1 + a1 \cdot pc) \cdot uCA_{PV}$$

By the Quasi-Steady State Approximation:

$$\frac{\mathrm{d}uCA_{PV}}{\mathrm{d}t} = 0$$

$$\frac{Q_{MA}}{SYS} \cdot (1 + a1 \cdot pc) \cdot uCA_{SYS} + klintabs_{uCA} \cdot lintabs_{bsv} \cdot uCA_{LINT} + kcolabs_{CA} \cdot colabs_{bsv} \cdot uCA_{COL} - \frac{Q_{PV}}{PV} \cdot (1 + a1 \cdot pc) \cdot uCA_{PV} = 0$$

Solve for $uCA_{PV}$:

$$uCA_{PV} = \frac{\frac{Q_{MA}}{SYS} \cdot (1 + a1 \cdot pc) \cdot uCA_{SYS} + klintabs_{uCA} \cdot lintabs_{bsv} \cdot uCA_{LINT} + kcolabs_{CA} \cdot colabs_{bsv} \cdot uCA_{COL}}{\frac{Q_{PV}}{PV} \cdot (1 + a1 \cdot pc)} \tag{1}$$

In a similar fashion the other portal vein bile acids can be calculated algebraically.

$$uCDCA_{PV} = \frac{\frac{Q_{MA}}{SYS} \cdot (1 + a1 \cdot pc) \cdot uCDCA_{SYS} + klintabs_{uCDCADCA} \cdot lintabs_{bsv} \cdot uCDCA_{LINT} + kcolabs_{CDCA} \cdot colabs_{bsv} \cdot uCDCA_{COL}}{\frac{Q_{PV}}{PV} \cdot (1 + a1 \cdot pc)} \tag{2}$$

$$uDCA_{PV} = \frac{\frac{Q_{MA}}{SYS} \cdot (1 + a1 \cdot pc) \cdot uDCA_{SYS} + klintabs_{uCDCADCA} \cdot lintabs_{bsv} \cdot uDCA_{LINT} + kcolabs_{DCA} \cdot colabs_{bsv} \cdot uDCA_{COL}}{\frac{Q_{PV}}{PV} \cdot (1 + a1 \cdot pc)} \tag{3}$$

##### Conjugated Bile Acids

$$tCA_{PV} = \frac{\frac{Q_{MA}}{SYS} \cdot (1 + a1 \cdot pc) \cdot tCA_{SYS} + klintabs_{tCA} \cdot lintabs_{bsv} \cdot tCA_{LINT}}{\frac{Q_{PV}}{PV} \cdot (1 + a1 \cdot pc)} \tag{4}$$

$$gCA_{PV} = \frac{\frac{Q_{MA}}{SYS} \cdot (1 + a1 \cdot pc) \cdot gCA_{SYS} + klintabs_{gCA} \cdot lintabs_{bsv} \cdot gCA_{LINT}}{\frac{Q_{PV}}{PV} \cdot (1 + a1 \cdot pc)} \tag{5}$$

$$tCDCA_{PV} = \frac{\frac{Q_{MA}}{SYS} \cdot (1 + a1 \cdot pc) \cdot tCDCA_{SYS} + klintabs_{tCDCA} \cdot lintabs_{bsv} \cdot tCDCA_{LINT}}{\frac{Q_{PV}}{PV} \cdot (1 + a1 \cdot pc)} \tag{6}$$

$$gCDCA_{PV} = \frac{\frac{Q_{MA}}{SYS} \cdot (1+a1 \cdot pc) \cdot gCDCA_{SYS} + kintabs_{tCDCA} \cdot uintabs_{bsv} \cdot gCDCA_{UINT} + klintabs_{gCDCA} \cdot lintabs_{bsv} \cdot gCDCA_{LINT}}{\frac{Q_{PV}}{PV} \cdot (1+a1 \cdot pc)} \tag{7}$$

$$tDCA_{PV} = \frac{\frac{Q_{MA}}{SYS} \cdot (1 + a1 \cdot pc) \cdot tDCA_{SYS} + klintabs_{tDCA} \cdot lintabs_{bsv} \cdot tDCA_{LINT}}{\frac{Q_{PV}}{PV} \cdot (1 + a1 \cdot pc)} \tag{8}$$

$$gDCA_{PV} = \frac{\frac{Q_{MA}}{SYS} \cdot (1+a1 \cdot pc) \cdot gDCA_{SYS} + kintabs_{gDCA} \cdot uintabs_{bsv} \cdot gDCA_{UINT} + klintabs_{gDCA} \cdot lintabs_{bsv} \cdot gDCA_{LINT}}{\frac{Q_{PV}}{PV} \cdot (1+a1 \cdot pc)} \tag{9}$$

### Sinusoidal Space

#### Unconjugated Bile Acids

$$\frac{\mathrm{d}uCA_{SIN}}{\mathrm{d}t} = \frac{Q_{PV}}{PV} \cdot (1 + a1 \cdot pc) \cdot uCA_{PV} + \frac{Q_{HA}}{SYS} \cdot (1 + a1 \cdot pc) \cdot uCA_{SYS} + kref \cdot uCA_L - \frac{Q_{HV}}{SIN} \cdot (1 + a1 \cdot pc) \cdot uCA_{SIN} - kupt_{uCA} \cdot hupt_{bsv} \cdot uCA_{SIN}$$

By the Quasi-Steady State Approximation:

$$\frac{\mathrm{d}uCA_{SIN}}{\mathrm{d}t} = 0$$

$$\frac{Q_{PV}}{PV} \cdot (1 + a1 \cdot pc) \cdot uCA_{PV} + \frac{Q_{HA}}{SYS} \cdot (1 + a1 \cdot pc) \cdot uCA_{SYS} + kref \cdot uCA_L - \frac{Q_{HV}}{SIN} \cdot (1 + a1 \cdot pc) \cdot uCA_{SIN} - kupt_{uCA} \cdot hupt_{bsv} \cdot uCA_{SIN} = 0$$

Solve for $uCA_{SIN}$:

$$uCA_{SIN} = \frac{\frac{Q_{PV}}{PV} \cdot (1 + a1 \cdot pc) \cdot uCA_{PV} + \frac{Q_{HA}}{SYS} \cdot (1 + a1 \cdot pc) \cdot uCA_{SYS} + kref \cdot uCA_L}{\frac{Q_{HV}}{SIN} \cdot (1 + a1 \cdot pc) + kupt_{uCA} \cdot hupt_{bsv}} \tag{10}$$

In a similar fashion the other sinusoidal space bile acids can be calculated algebraically.

$$uCDCA_{SIN} = \frac{\frac{Q_{PV}}{PV} \cdot (1 + a1 \cdot pc) \cdot uCDCA_{PV} + \frac{Q_{HA}}{SYS} \cdot (1 + a1 \cdot pc) \cdot uCDCA_{SYS} + kref \cdot uCDCA_L}{\frac{Q_{HV}}{SIN} \cdot (1 + a1 \cdot pc) + kupt_{uCDCA} \cdot hupt_{bsv}} \tag{11}$$

$$uDCA_{SIN} = \frac{\frac{Q_{PV}}{PV} \cdot (1 + a1 \cdot pc) \cdot uDCA_{PV} + \frac{Q_{HA}}{SYS} \cdot (1 + a1 \cdot pc) \cdot uDCA_{SYS} + kref \cdot uDCA_L}{\frac{Q_{HV}}{SIN} \cdot (1 + a1 \cdot pc) + kupt_{uDCA} \cdot hupt_{bsv}} \tag{12}$$

#### Conjugated Bile Acids

$$tCA_{SIN} = \frac{\frac{Q_{PV}}{PV} \cdot (1 + a1 \cdot pc) \cdot tCA_{PV} + \frac{Q_{HA}}{SYS} \cdot (1 + a1 \cdot pc) \cdot tCA_{SYS} + kref \cdot tCA_L}{\frac{Q_{HV}}{SIN} \cdot (1 + a1 \cdot pc) + kupt_{tCA} \cdot hupt_{bsv}} \tag{13}$$

$$gCA_{SIN} = \frac{\frac{Q_{PV}}{PV} \cdot (1 + a1 \cdot pc) \cdot gCA_{PV} + \frac{Q_{HA}}{SYS} \cdot (1 + a1 \cdot pc) \cdot gCA_{SYS} + kref \cdot gCA_L}{\frac{Q_{HV}}{SIN} \cdot (1 + a1 \cdot pc) + kupt_{gCA} \cdot hupt_{bsv}} \tag{14}$$

$$tCDCA_{SIN} = \frac{\frac{Q_{PV}}{PV} \cdot (1 + a1 \cdot pc) \cdot tCDCA_{PV} + \frac{Q_{HA}}{SYS} \cdot (1 + a1 \cdot pc) \cdot tCDCA_{SYS} + kref \cdot tCDCA_L}{\frac{Q_{HV}}{SIN} \cdot (1 + a1 \cdot pc) + kupt_{tCDCA} \cdot hupt_{bsv}} \tag{15}$$

$$gCDCA_{SIN} = \frac{\frac{Q_{PV}}{PV} \cdot (1 + a1 \cdot pc) \cdot gCDCA_{PV} + \frac{Q_{HA}}{SYS} \cdot (1 + a1 \cdot pc) \cdot gCDCA_{SYS} + kref \cdot gCDCA_L}{\frac{Q_{HV}}{SIN} \cdot (1 + a1 \cdot pc) + kupt_{gCDCA} \cdot hupt_{bsv}} \tag{16}$$

$$tDCA_{SIN} = \frac{\frac{Q_{PV}}{PV} \cdot (1 + a1 \cdot pc) \cdot tDCA_{PV} + \frac{Q_{HA}}{SYS} \cdot (1 + a1 \cdot pc) \cdot tDCA_{SYS} + kref \cdot tDCA_L}{\frac{Q_{HV}}{SIN} \cdot (1 + a1 \cdot pc) + kupt_{tDCA} \cdot hupt_{bsv}} \tag{17}$$

$$gDCA_{SIN} = \frac{\frac{Q_{PV}}{PV} \cdot (1 + a1 \cdot pc) \cdot gDCA_{PV} + \frac{Q_{HA}}{SYS} \cdot (1 + a1 \cdot pc) \cdot gDCA_{SYS} + kref \cdot gDCA_L}{\frac{Q_{HV}}{SIN} \cdot (1 + a1 \cdot pc) + kupt_{gDCA} \cdot hupt_{bsv}} \tag{18}$$
