## Supplementary Figures for "Temporal blood sampling improves parameter recovery in a model of enterohepatic bile acid circulation using surrogate-accelerated Bayesian inference"

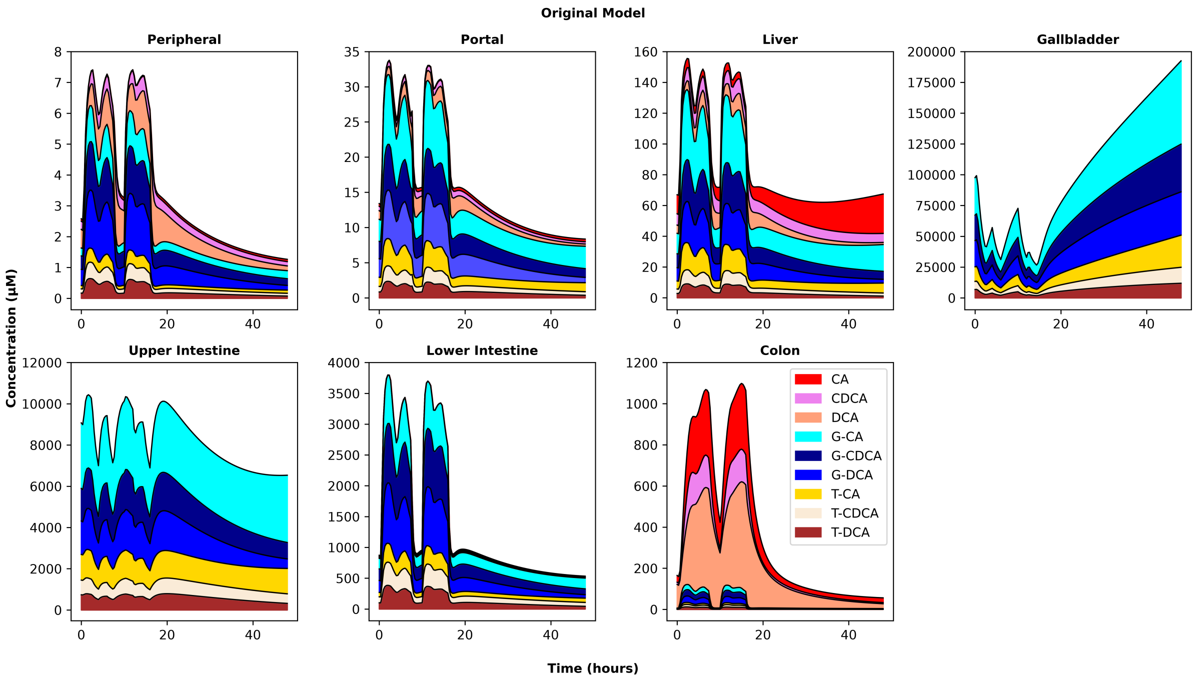


Figure 1: **Predicted bile acid dynamics in the original Voronova *et al.* model.** Concentrations of nine bile acid species across seven compartments (peripheral blood, portal blood, liver, gallbladder, upper intestine, lower intestine, and colon) over a 24-hour limit cycle followed by 24 hours of fasting. Abbreviations: *CA, cholic acid; CDCA, chenodeoxycholic acid; DCA, deoxycholic acid; G-, glycine-conjugated; T-, taurine-conjugated.*

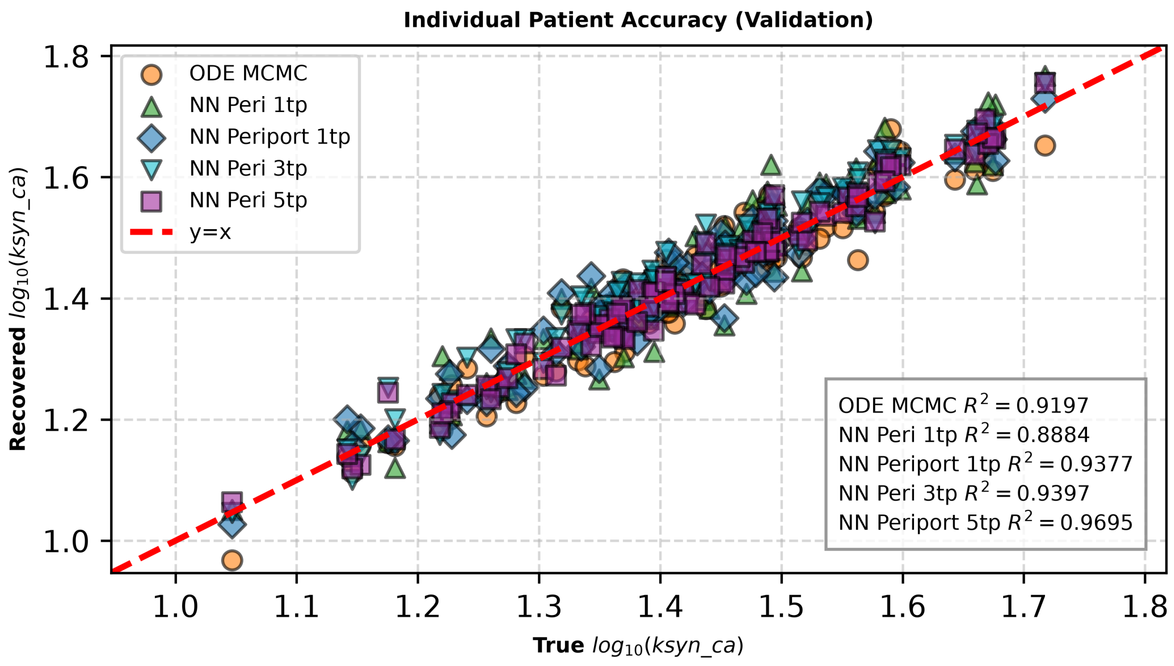


Figure 2: **Neural network surrogates reproduce single parameter Bayesian inference across all sampling protocols.** Parity plots showing the posterior means of $\log_{10}(\texttt{ksyn\_ca})$ recovered by the QSSA-reduced ODE model and the four NN-derived surrogates (Peri 1tp, Periport 1tp, Peri 3tp, Peri 5tp) against the true virtual patient values. All protocols achieve strong coefficients of determination ($R^2 > 0.88$), confirming that the surrogates do not introduce bias and that each sampling design contains sufficient information to identify ksyn_ca in isolation, validating the surrogate-accelerated pipeline before extending to five-parameter inference. Abbreviations: *MCMC, Markov chain Monte Carlo; NN, neural network; ODE, ordinary differential equation; QSSA, quasi-steady state approximation.*

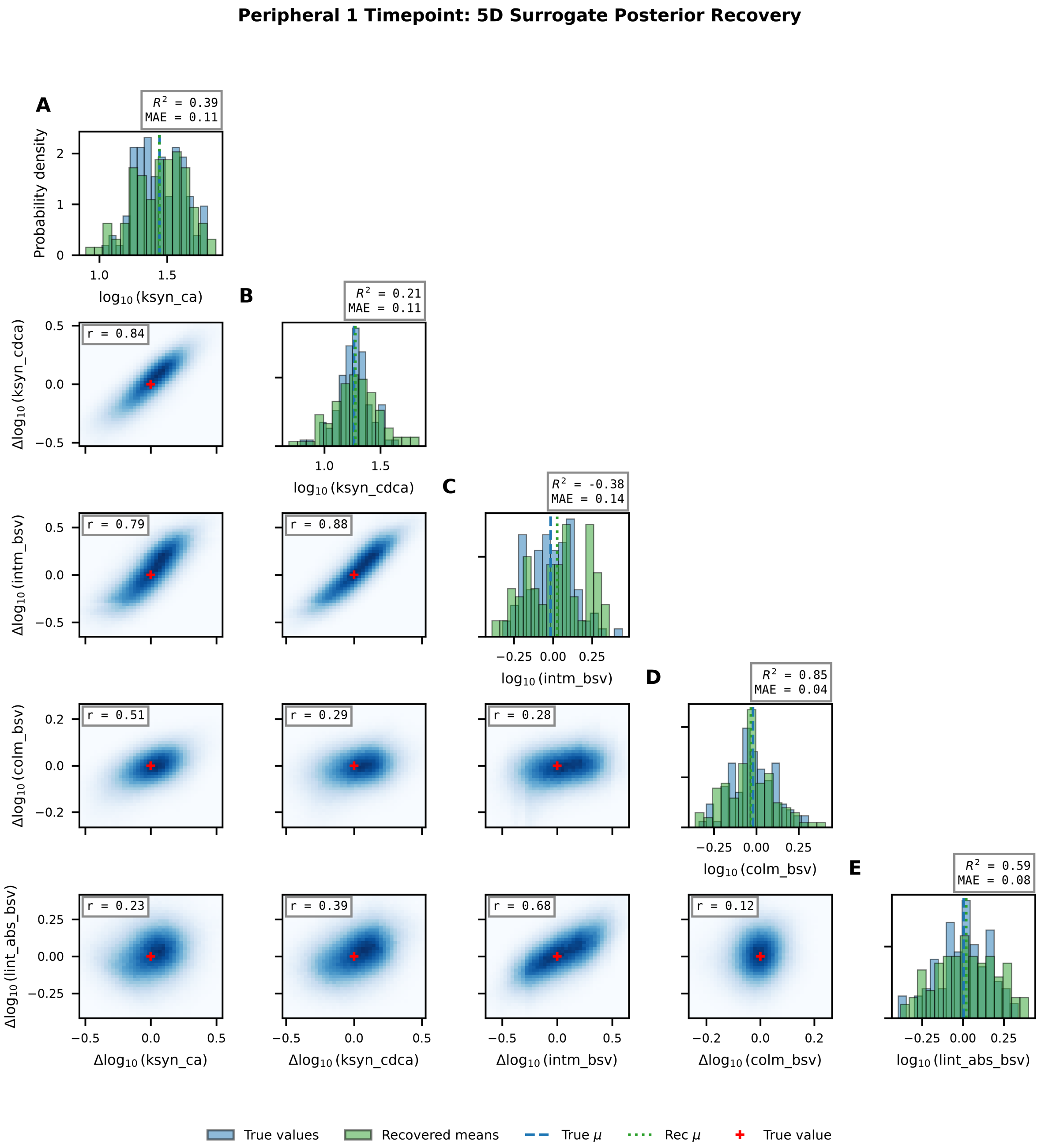


Figure 3: **Full five-parameter joint posterior distributions under single time point peripheral blood sampling (Peri 1tp)** (A-E) Marginal histograms of true (blue) and recovered (green) $\log_{10}$ parameter values for $N = 100$ virtual patients, with $R^2$ and MAE annotated. Triangle plot of re-centred joint posteriors ($\Delta \log_{10}$ = sample − true value) pooled across all patients, with median within-patient correlations annotated. This protocol exhibits severe practical non-identifiability, particularly for `intm_bsv` ($R^2 = -0.40$). *Abbreviations: colm_bsv, colonic motility scalar; intm_bsv, intestinal motility scalar; ksyn_ca, hepatic synthesis rate of cholic acid; ksyn_cdca, hepatic synthesis rate of chenodeoxycholic acid; lint_abs_bsv, lower intestinal absorption scalar; MAE, mean absolute error; $R^2$, coefficient of determination*

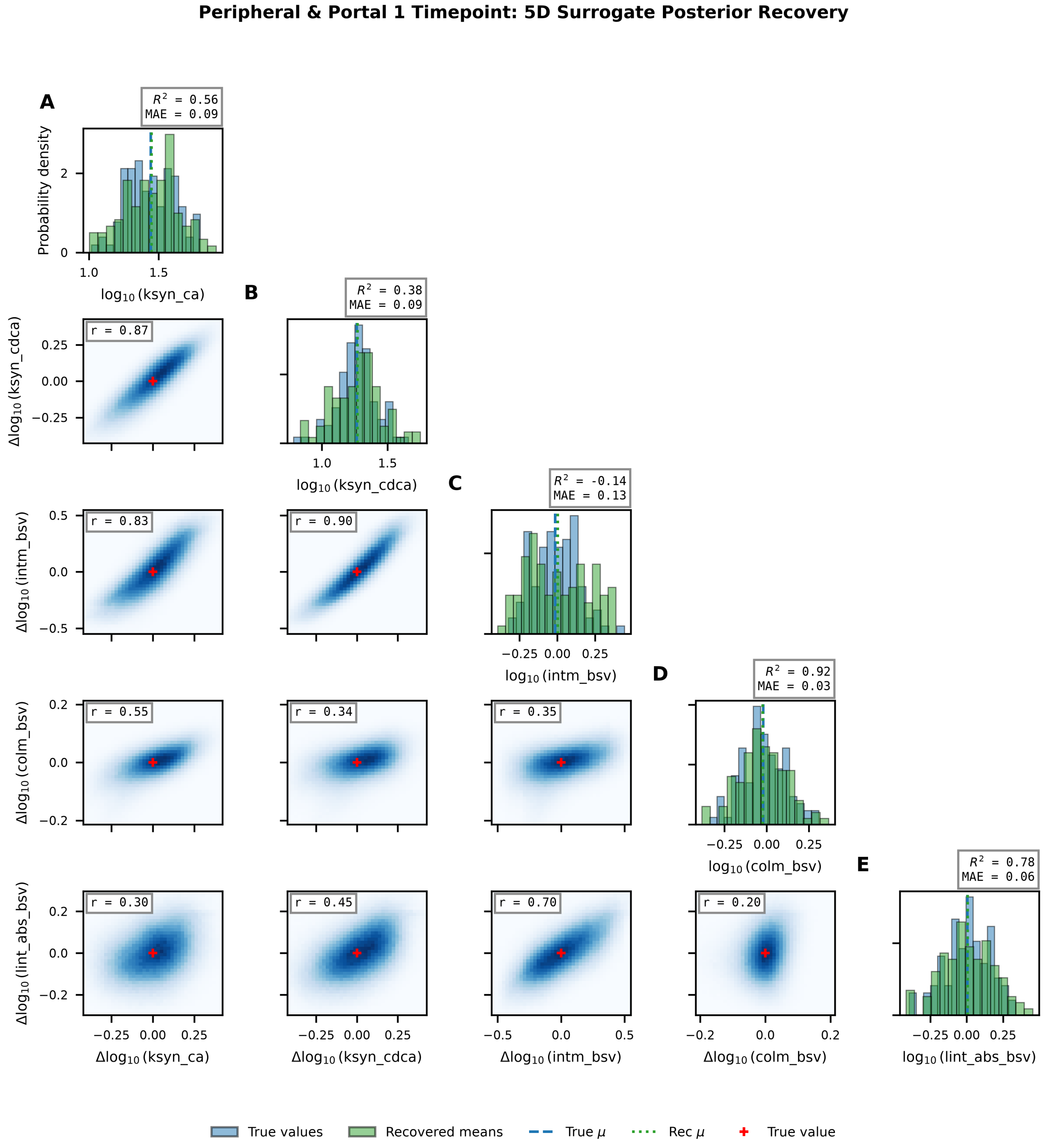


Figure 4: **Full five-parameter joint posterior distributions under single time point peripheral and portal blood sampling (Periport 1tp)** Layout as in S4 Fig. Augmenting peripheral measurements with portal vein concentrations improves recovery of `lint_abs_bsv` ($R^2 = 0.78$) but fails to resolve `intm_bsv` ($R^2 = -0.13$), demonstrating that spatial diversity alone cannot break the synthesis-transit degeneracy. *Abbreviations: colm_bsv, colonic motility scalar; intm_bsv, intestinal motility scalar; ksyn_ca, hepatic synthesis rate of cholic acid; ksyn_cdca, hepatic synthesis rate of chenodeoxycholic acid; lint_abs_bsv, lower intestinal absorption scalar; MAE, mean absolute error;* $R^2$*, coefficient of determination*

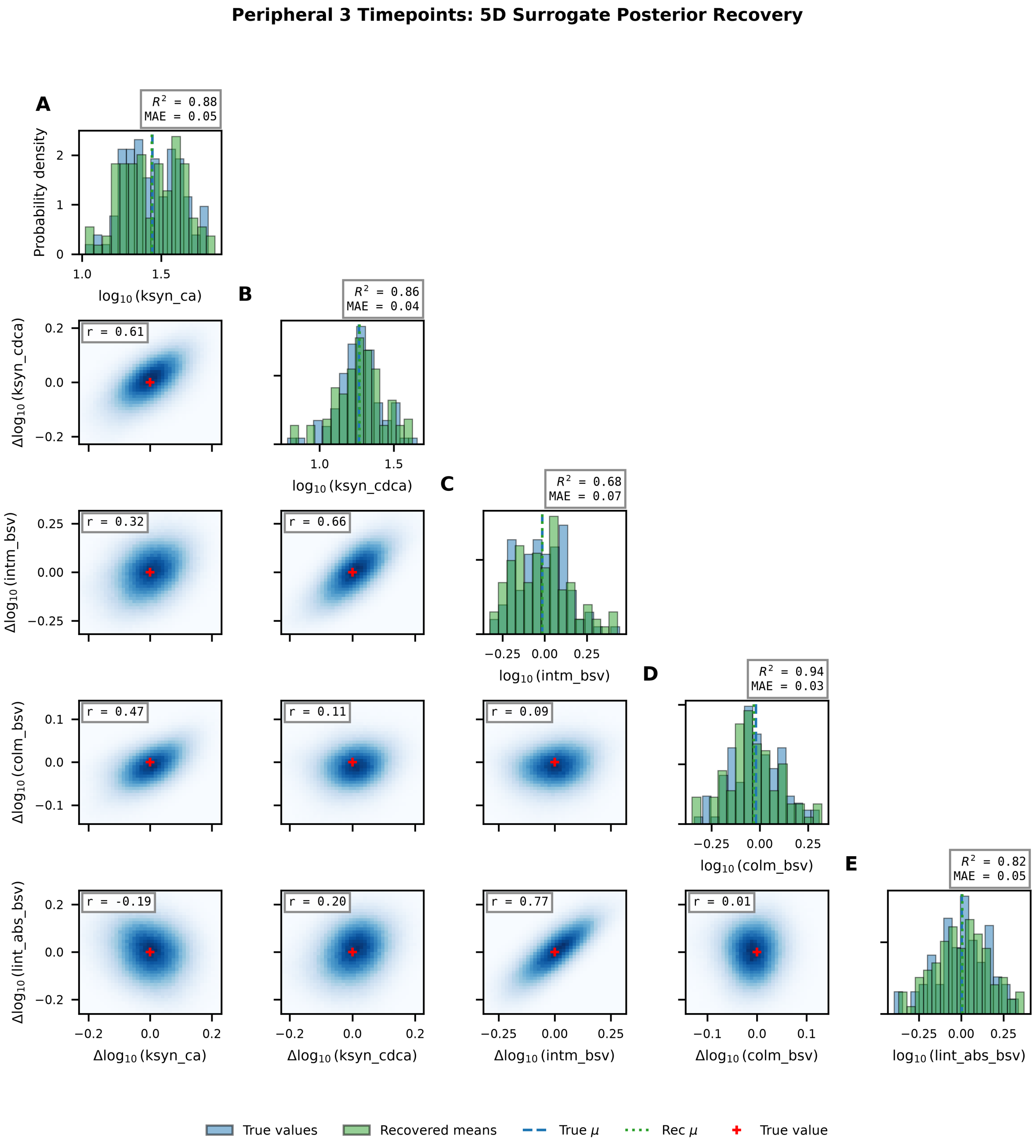


Figure 5: **Full five-parameter joint posterior distributions under peripheral blood sampling at three time points (Peri 3tp).** Layout as in S4 Fig. Multi-timepoint sampling resolves all five parameters with $R^2 \geq 0.68$, breaking the primary synthesis-transit degeneracy (median $r = 0.33$). *Abbreviations: colm_bsv, colonic motility scalar; intm_bsv, intestinal motility scalar; ksyn_ca, hepatic synthesis rate of cholic acid; ksyn_cdca, hepatic synthesis rate of chenodeoxycholic acid; lint_abs_bsv, lower intestinal absorption scalar; MAE, mean absolute error;* $R^2$*, coefficient of determination*

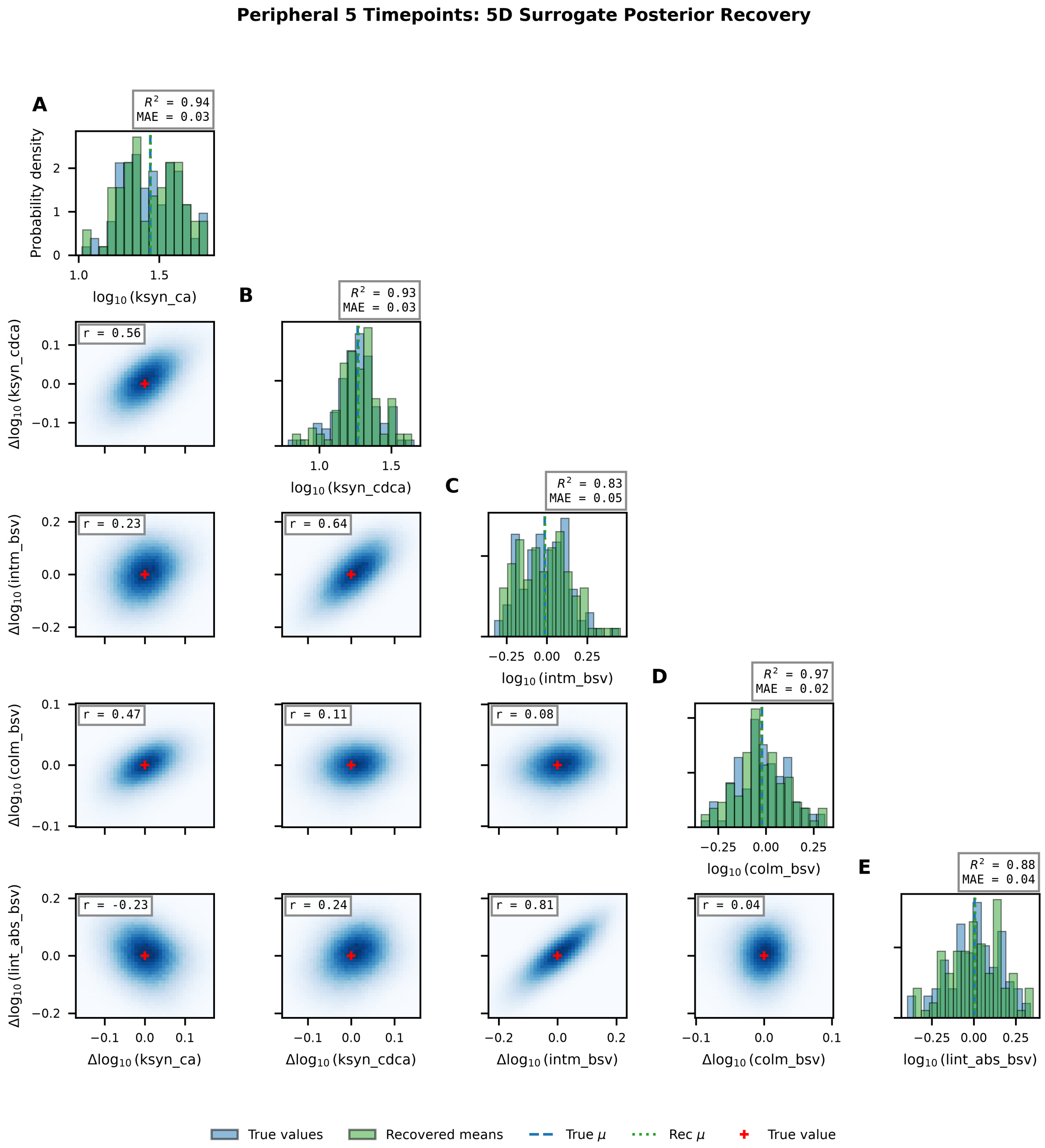


Figure 6: **Full five-parameter joint posterior distributions under peripheral blood sampling at five time points (Peri 5tp).** Layout as in S4 Fig. The sampling protocol achieves $R^2 \geq 0.83$ for all parameters and further reduces the synthesis-transit correlation to $r = 0.22$. *Abbreviations: colm_bsv, colonic motility scalar; intm_bsv, intestinal motility scalar; ksyn_ca, hepatic synthesis rate of cholic acid; ksyn_cdca, hepatic synthesis rate of chenodeoxycholic acid; lint_abs_bsv, lower intestinal absorption scalar; MAE, mean absolute error;* $R^2$*, coefficient of determination*

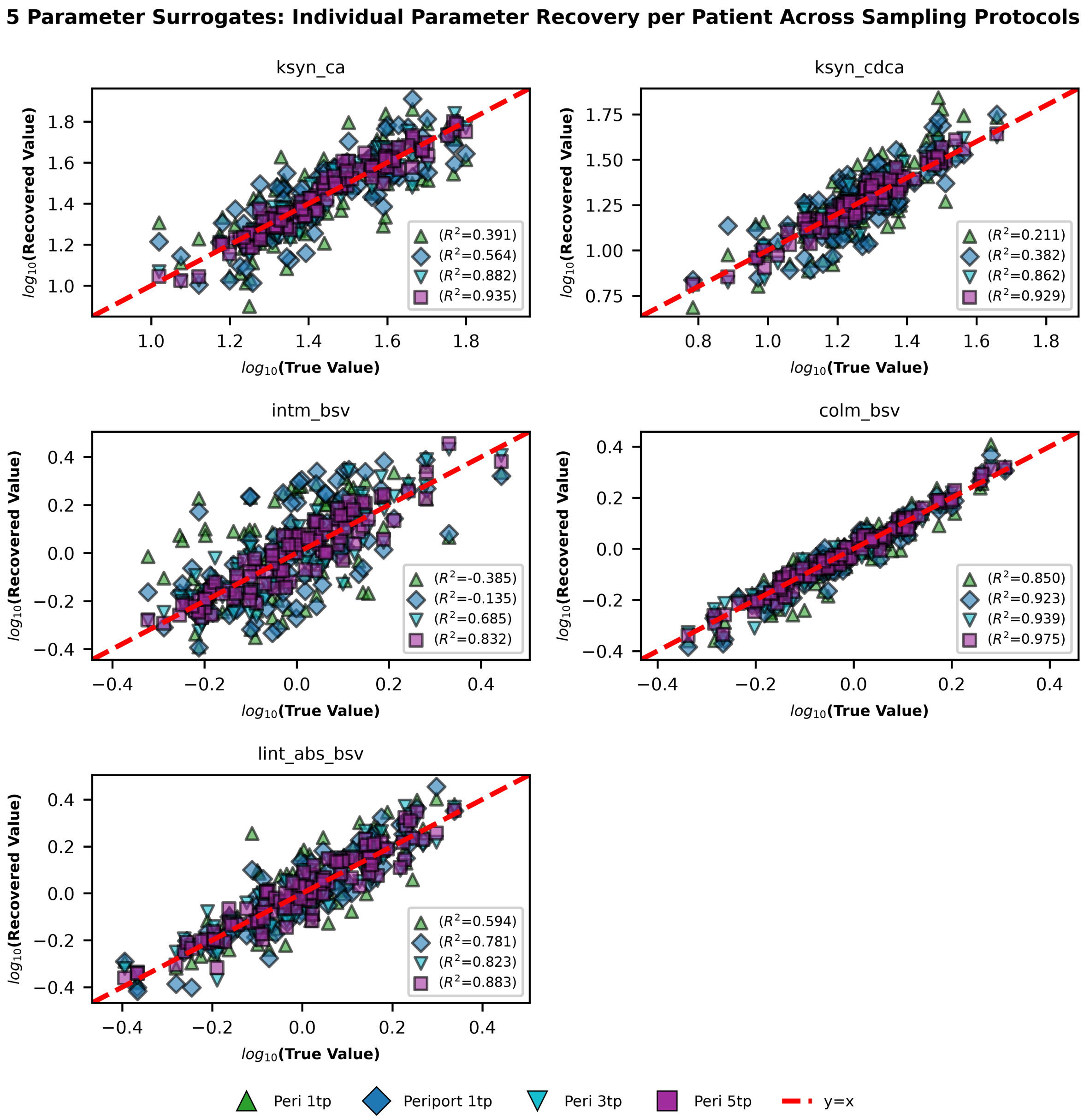


Figure 7: **Individual patient recovery accuracy for five parameter inference across all sampling protocols.** Parity plots showing the posterior means of each parameter recovered by the NN surrogates for each sampling protocol against the true virtual patient values. Each panel corresponds to one parameter, with all four protocols overlaid. The Peri 5tp protocol (purple squares) achieves the tightest clustering around the $y = x$ line for all parameters, while Peri 1tp (green triangles) shows the greatest scatter, particularly for `intm_bsv`. *Abbreviations: colm_bsv, colonic motility scalar; intm_bsv, intestinal motility scalar; ksyn_ca, hepatic synthesis rate of cholic acid; ksyn_cdca, hepatic synthesis rate of chenodeoxycholic acid; lint_abs_bsv, lower intestinal absorption scalar; Peri, peripheral; Periport, peripheral + portal;* $R^2$*, coefficient of determination; tp, time point.*

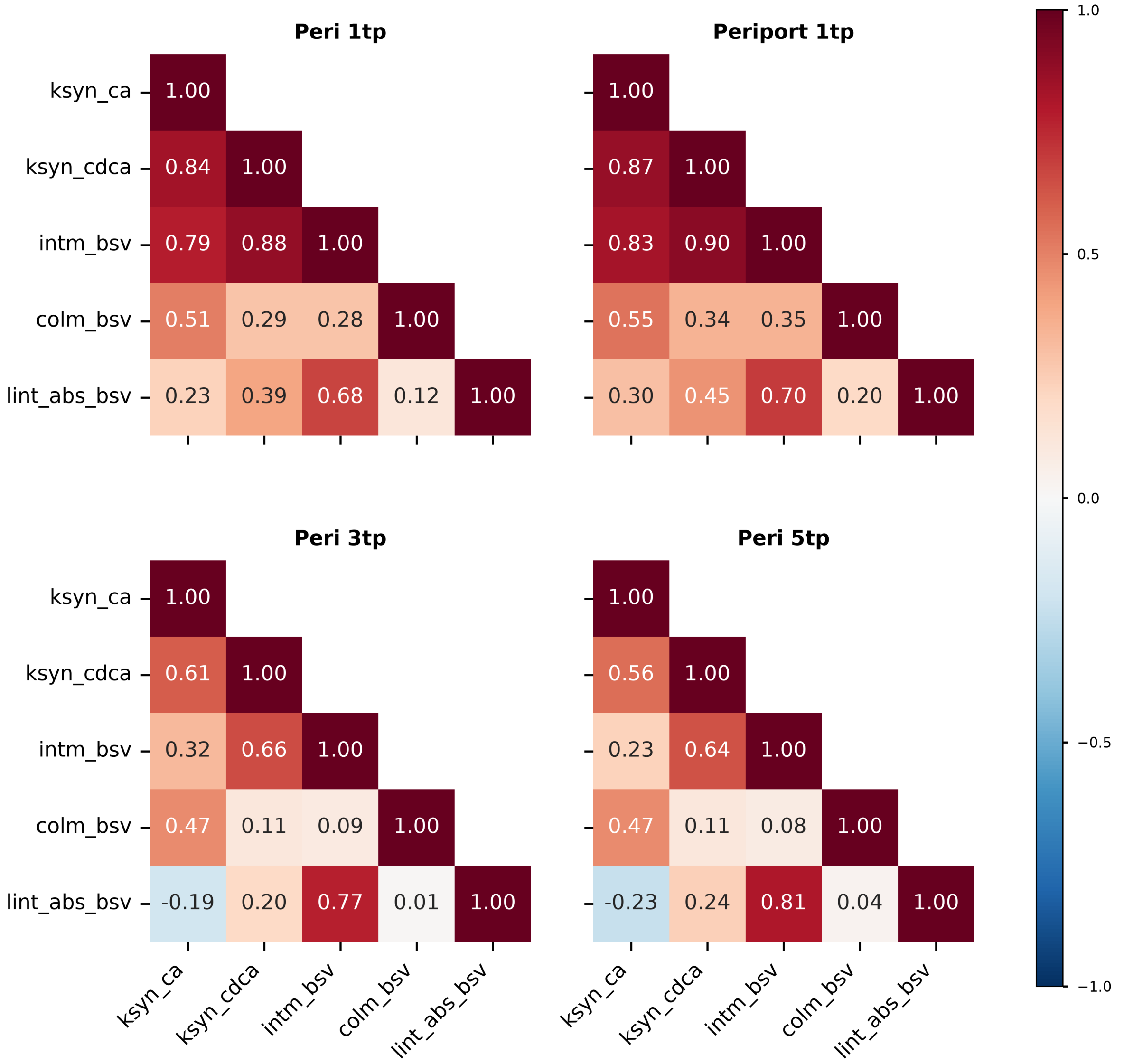


Figure 8: **Median within-patient posterior correlation matrices across all sampling protocols.** Heatmaps showing the median Pearson correlation coefficient ($r$) between all parameter pairs, computed across the $N = 100$ virtual patients. Single-timepoint protocols (Peri 1tp, Periport 1tp) exhibit strong positive correlations between synthesis rates and intestinal motility ($r \approx 0.79$–$0.83$), reflecting the synthesis–transit degeneracy. Multi-timepoint protocols (Peri 3tp, Peri 5tp) progressively decouple these parameters, with the Peri 5tp protocol achieving near-zero correlations for most pairs. *Abbreviations: colm_bsv, colonic motility scalar; intm_bsv, intestinal motility scalar; ksyn_ca, hepatic synthesis rate of cholic acid; ksyn_cdca, hepatic synthesis rate of chenodeoxycholic acid; lint_abs_bsv, lower intestinal absorption scalar; Peri, peripheral; Periport, peripheral + portal; tp, time point.*

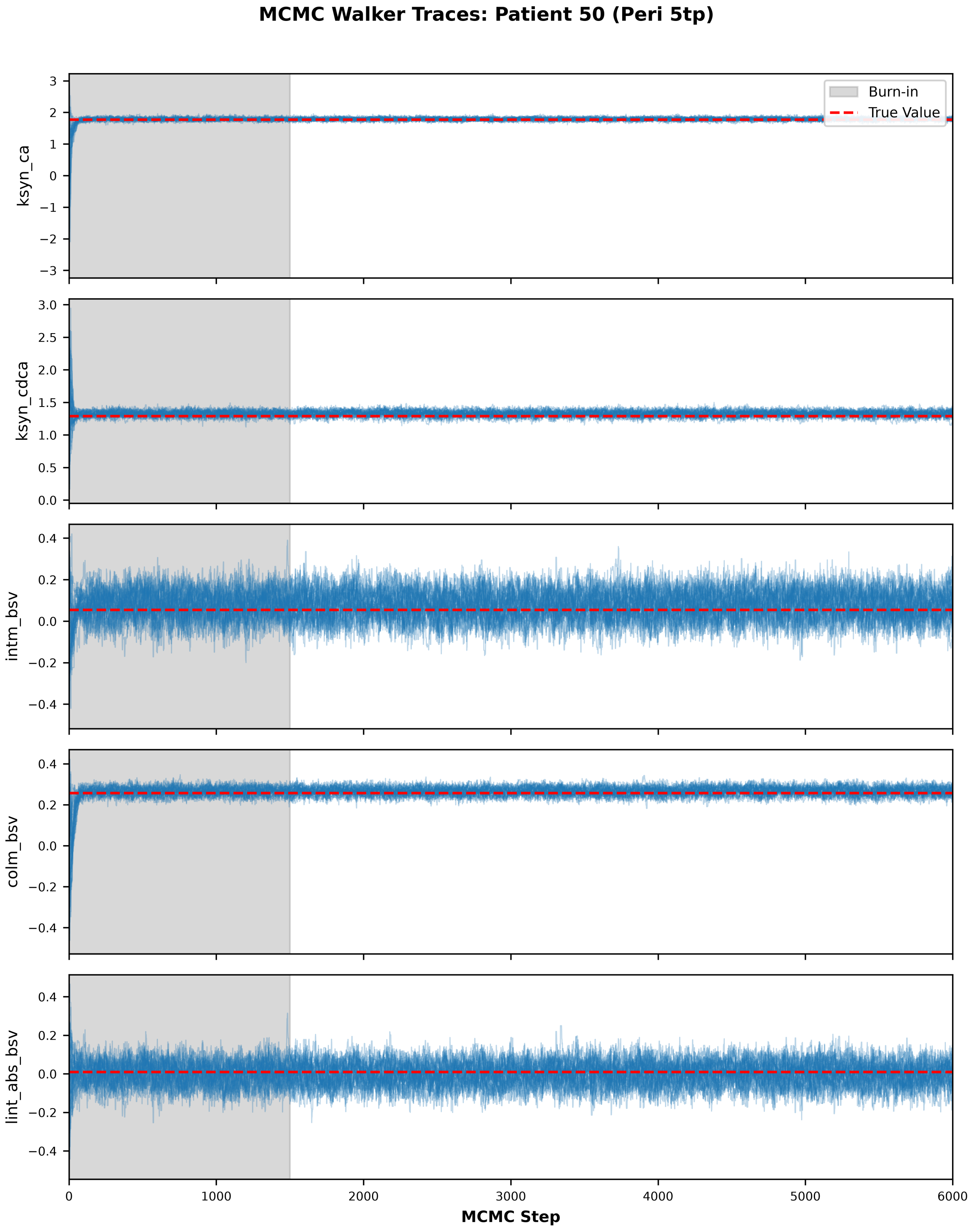


Figure 9: **MCMC convergence for the Peri 5tp protocol.** Trace plots showing the evolution of parameter samples across the 6,000 MCMC steps for a representative virtual patient. The burn-in period (first 1,500 steps) is shaded in grey.
